# Exploring Neural Heterogeneity in Inattention and Hyperactivity

**DOI:** 10.1101/2022.07.26.501508

**Authors:** Natalia Zdorovtsova, Jonathan Jones, Danyal Akarca, Elia Benhamou, The CALM Team, Duncan E. Astle

## Abstract

Inattention and hyperactivity are cardinal symptoms of Attention Deficit Hyperactivity Disorder (ADHD). These characteristics have also been observed across a range of other neurodevelopmental conditions, such as autism and dyspraxia, suggesting that they might best be studied across diagnostic categories. Here, we evaluated the associations between inattention and hyperactivity behaviours and features of the structural brain network (connectome) in a large transdiagnostic sample of children (Centre for Attention, Learning, and Memory; n = 383). In our sample, we found that a single latent factor explains 77.6% of variance in scores across multiple questionnaires measuring inattention and hyperactivity. Partial Least-Squares (PLS) regression revealed that variability in this latent factor could not be explained by a linear component representing nodewise properties of connectomes. We then investigated the type and extent of neural heterogeneity in a subset of our sample with clinically-elevated levels of inattention and hyperactivity. Multidimensional scaling combined with k-means clustering revealed two neural subtypes in children with elevated levels of inattention and hyperactivity (n = 232), differentiated primarily by nodal communicability—a measure which demarcates the extent to which neural signals propagate through specific brain regions. These different clusters had indistinguishable behavioural profiles, which included high levels of inattention and hyperactivity. However, one of the clusters scored higher on multiple cognitive assessment measures of executive function. We conclude that inattention and hyperactivity are so common in children with neurodevelopmental difficulties because they emerge from multiple different trajectories of brain development. In our own data, we can identify two of these possible trajectories, which are reflected by measures of structural brain network topology and cognition.

**Research Highlights:**

- We investigated variability in structural brain network organisation and its relationship with cognition and behaviour in a sample of 383 children.
- We did not find linear components of brain structure that explained continuous variations in inattention and hyperactivity across this heterogeneous sample.
- Following this, we explored different attributes of brain organisation in children with particularly elevated levels of inattention and hyperactivity (n = 232).
- Among highly inattentive and hyperactive children, we found two profiles of structural brain organisation (‘neurotypes’), which were differentiated primarily by the communicability of nodes in frontal and occipital brain areas.
- These subgroups did not differ on additional measures of behaviour. However, the lower-nodal-communicability group demonstrated weaker performance on cognitive assessments of executive function and visuospatial processing.
- We discuss the implications that these findings have for our understanding of variability in neurodevelopmental difficulties and related conditions, such as ADHD

## Introduction

### A Transdiagnostic Approach

Neurodevelopmental conditions like ADHD, autism, and dyslexia have traditionally been regarded as fundamentally-distinct diagnostic categories in both research and clinical practice. As a result, past research in developmental psychology, child psychiatry and neuroscience has prioritised the comparison of highly-selective clinical samples with ‘typically-developing’ controls. Approaches to sampling, experimental design, and data analysis have been shaped by the assumption of underlying uniformity within, and distinctiveness between, diagnostic categories (Coghill & Sonuga-Barke, 2012; Dalgleish et al., 2020; Astle et al., 2022). This creates an organisational framework for studying cognitive, behavioural, and neural differences between groups. However, this case-control framework struggles to accommodate the reality of neurodevelopmental diversity, and current diagnostic labels do not actually reflect self-contained developmental phenotypes.

The degree and co-occurrence of clinical features across individuals experiencing neurodevelopmental difficulties is highly heterogeneous. There is significant cognitive and behavioural overlap between individuals who supposedly have different disorders (e.g. Jette & Geschwind, 2014; Jacob et al., 2019; Kofler et al., 2019). Certain cognitive and behavioural symptoms—like emotional dysregulation, inattention, and differences in sensory perception—are also commonly observed across multiple neurodevelopmental conditions, and in those with no formal diagnosis (Lau-Zhu et al., 2019; Krakowski et al., 2020; Dellapiazza et al., 2020). Additionally, ADHD and autism have diagnostic co-occurrence rates of up to 70% (Reiersen & Todd, 2008; Joshi et al., 2017). Due to the heterogeneity of symptoms across neurodevelopmental conditions, the traditional dichotomisation of ‘typically-developing’ and ‘atypically-developing’ or ‘Diagnosis A’ versus ‘Diagnosis B’ has been increasingly challenged (see Fusar-Poli et al., 2019 and Astle & Fletcher-Watson, 2020, for review). Put simply, a rigid categorisation of individuals according to conventional diagnostic labels fails to adequately capture the similarities and differences that exist between individuals.

Some research groups have come to favour a transdiagnostic approach as an alternative to the conventional case-control design (e.g. Vanden Bussche et al., 2017; Holmes et al., 2019; Shephard et al., 2019; Gui et al., 2020; Hayiou-Thomas et al., 2021). Transdiagnostic approaches place less priority on the diagnostic categorisation of conditions, instead focusing on variations in the dimensionality of traits, often with a broader representation of co-occurring conditions within participant samples (e.g. Cuthbert, 2014; Newby et al., 2015; Talkovsky et al., 2017; Bathelt et al., 2018; Dalgleish et al., 2020; Siugzdaite et al., 2020). Advocates for transdiagnostic approaches endeavour to build an analytic and conceptual framework that accommodates heterogeneity within, and between, neurodevelopmental conditions. One aim is to reconceptualise neurodevelopmental conditions as intrinsically related, not only in terms of their behavioural attributes, but also potentially the neural and cognitive mechanisms which underpin them (Forest, 2014).

Inattention and hyperactivity are hallmark symptoms of ADHD, but they nonetheless occur across the entire population along a continuous spectrum (see Nigg, 2013, for review). Unusually high levels of inattention and hyperactivity are associated with multiple neurodevelopmental conditions, including ADHD, Autism, and Intellectual Disability (McClain et al., 2017; Krakowski et al., 2020). This suggests that inattention and hyperactivity, as features of behaviour, do not strongly delineate different neurodevelopmental conditions. The purpose of the current study was to explore inattention and hyperactivity as transdiagnostic symptom dimensions, and to understand how these dimensions relate to variability in brain organisation.

### Brain Connectomics

In recent decades, a variety of fields have embraced the study of complex networks (Sporns et al., 2005; Betzel, 2022). Characterising the topological properties of complex networks allows us to summarise network architecture across multiple scales and levels of interaction, rather than being limited to narrow, specific regions of the brain assumed to operate independently (Costa et al., 2007). The human brain is a complex system comprised of manifold intricate connections between neurons, optimised to maximise efficient information transfer whilst minimising energy usage (Bullmore and Sporns, 2012). The connection matrix of the human brain (hereafter ‘connectome’) provides a useful model of its anatomical structure and function (Sporns et al., 2005).

Graph theory, a branch of mathematics, enables the quantitative modelling of network characteristics. This provides a formal framework for characterising structural and functional connectomes (Trudeau, 2013). A graph’s information is typically contained within an adjacency matrix, which represents connections (‘edges’) between entities (‘nodes’). Edges can be weighted (continuous) or binary (present vs. absent), and they can represent directed or undirected relationships of connectivity between nodes. Macroscopic-level connectomics characterises brain regions, which are defined using a range of possible brain atlases, as nodes. Edges represent patterns and degrees of structural, functional, and effective (causal) relationships between those nodes (Rubinov & Sporns, 2010). Graph-theoretic measures at the level of nodes, commonly named centrality indices, contain information that can provide insight into the role of a specific brain region in the overall organisation of the brain (Buckholtz & Meyer-Lindenberg, 2012; van Essen & Barch, 2015).

The white matter connectome is thought to constrain the intertwined communication dynamics of the brain, facilitating function (Avena-Koenigsberger et al., 2017; Suárez et al., 2021). Variations in this communication network may coincide with differences in cognition and behaviour, in line with empirical observations in children (e.g. Mareva et al., 2019; Siugzdaite et al., 2020). In the current study, we explored how node-level measures in the structural brain network relate to variations in hyperactivity- and inattention-related behaviours. We tested how the number of connections (termed *degree*), triplets of connectivity (termed *clustering coefficients*) and their local patterning of communication (termed *communicability*) for different brain regions may vary according to a young person’s levels of inattention and hyperactivity. We used these brain network measures as proxies for network architecture, reduced to the regional level, to enable appropriate statistical inferences. We chose these measures because they are capable of describing various attributes of neural organisation—such as the formation of structural modules around nodes with high degree and clustering coefficient values, and the extent to which these nodes may facilitate information flow across the brain.

### Structural Brain Differences in Neurodevelopmental Conditions

The study of structural brain differences in those with neurodevelopmental conditions has existed at the core of human developmental psychology and neuroscience since the emergence of modern neuroimaging techniques. Certain features of brain structure and function have been linked to individual differences in behaviour and cognition in childhood (e.g. Astle et al., 2019; see Gilmore et al., 2018 and Ziegler et al., 2020 for review). These findings come mostly from studies of neurodevelopmental conditions like ADHD, autism, and dyslexia, which have attempted to draw contrasts between these groups and so-called typically-developing children (Griffiths et al., 2016; Mizuno et al., 2019).

To provide some brief examples from this literature: several studies report brain structure differences in autistic individuals, including grey matter reductions and increased gyrification in sensorimotor and default-mode networks (Pereira et al., 2018), reduced interhemispheric connectivity (Bos et al., 2015), and differences in long-range white matter connectivity (Wilkinson et al., 2016). In the case of both ADHD and autism, studies have shown similar patterns of increased brain network modularity, coupled with a global decrease in long-range connections (Kern et al., 2015), as well as white matter differences in frontal and limbic brain networks (Connaughton et al., 2022). Structural connectivity has also been studied in relation to specific behavioural traits associated with ADHD. Most relevant here, Konrad et al. (2010) found that inattention and impulsivity are related to alterations in fronto-striatal circuitry, which persists into adulthood.

More recent approaches to studying neurodevelopmental differences in structural connectivity have probed the nature of diversity as a feature of human brain network configurations. For instance, Akarca et al. (2021) used generative modelling to simulate the wiring properties of children at heightened risk for divergent neurodevelopmental outcomes. This analysis, without putting particular priority on any specific neurodevelopmental condition, revealed that the structural diversity we see in the macroscopic brain networks of children can be explained by small, constrained variations in generative parameter conditions. In other words, human brain development is not optimised to exhibit exactly the same wiring negotiations between regions, across individuals; rather, nature permits for structural differences across individuals, so long as some level of gross functionality is achieved. Diversity in the structure of human brains may be an adaptive or stochastic fundamental feature, not a flaw, of evolutionary and developmental processes. It is therefore valuable to investigate what, if anything, characterises structural brain differences in those experiencing impactful neurodevelopmental differences, such as heightened inattention and hyperactivity.

### The Current Study

We used data taken from a transdiagnostic cohort called the Centre for Attention, Learning and Memory (CALM) to test how properties of structural connectomes relate to variations in inattention and hyperactivity behavioural traits. The members of this cohort are recruited via referral from children’s specialist services on the basis of ongoing difficulties in cognition, learning or behaviour (Holmes et al., 2019). Cohort members can have single, multiple, or no formal diagnostic label(s).

We attempted to address one overarching question: how do properties of structural brain connectivity relate to variations along the transdiagnostic continua of inattention and hyperactivity within this intentionally-heterogeneous sample? We take two approaches to addressing this question. First, we test whether variability in inattention and hyperactivity within the sample relates to continuous variations in node-wise properties of each individual’s connectomes. Second, we test whether there are subgroups, with differently-organised connectomes, that experience inattention and hyperactivity.

## Methods and Materials

### Participants

The CALM sample was recruited on the basis of ongoing problems in attention, learning and memory reported by professionals working in schools or specialist children’s community services. Exclusion criteria for referrals were a known history of brain injury, significant or severe known problems in vision or hearing that were uncorrected, and not having an adequate level of English proficiency to complete the assessments. This study was approved by the local NHS research ethics committee (Reference: 13/EE/0157). Children attending the research clinic completed a cognitive test battery administered over approximately 4 hours, which covered a broad range of behavioural and cognitive attributes. Their parents also completed a set of standardised behavioural questionnaires assessing communication, inattention, hyperactivity, executive functioning and aspects of social functioning (details of the full protocol can be found in Holmes et al., 2019). Each child also underwent T1 structural MRI scanning at the MRC Cognition and Brain Sciences Unit. Several steps were taken to ensure good MRI data quality and minimize potential biases of participant movement, which has previously been found to affect statistical comparisons, particularly in hyperactive individuals (Yendiki et al., 2014). First, children were instructed to lie still and were trained to do so in a realistic mock scanner prior to the actual scan. Second, scans that showed a displacement of >3 mm within the diffusion-weighted sequence were excluded. We also used mean framewise displacement as a control regressor in our analyses across participants.

The main CALM cohort, from which we studied a subsample, includes n = 799 referred children and young people, alongside an additional n = 158 unreferred children and young people from the same schools. These additional unreferred participants were recruited as part of the cohort to ensure that the overall cohort captured the full range of cognitive and behavioural profiles in the wider population. At the time of our analysis, of the overall cohort (n = 957), 383 had complete and high-quality diffusion tensor imaging (DTI) data alongside cognitive and behavioural data (mean age = 10.33 years, SD = 2.27; 65.9% male, 33.1% female, 1% unspecified sex). In the current study, we analysed data from these 383 individuals.

### Cognitive Assessments and Behavioural Questionnaires

#### Behavioural Questionnaires

The Conners 3 - Parent Rating Scale Short Form was used to assess symptoms related to ADHD (Conners Questionnaire; Conners, 2008). Scores on these items form six subscales consisting of Inattention, Hyperactivity/Impulsivity, Learning Problems, Executive Function, Aggression, and Peer Relations. The sum of raw scores on each subscale was converted to a T-score (M = 50, SD = 10).

The Behavior Rating Inventory of Executive Function questionnaire was completed by parents/carers (BRIEF, Gioia et al., 2000). T-scores (M = 50, SD = 10) were derived for eight subscales: Inhibit, Shift, Emotional control, Initiate, Working memory, Planning, Organisation and Monitor. Three composite scores were also derived: Metacognition, Behaviour Regulation and Global Executive Function.

The Strengths and Difficulties Questionnaire asked the parent/carer to rate 25 items measuring Emotional Symptoms, Conduct Problems, Hyperactivity/Inattention, Peer Relationship Problems and Prosocial Behaviour based on their child’s behaviour in the six months prior to assessment (SDQ; Goodman, 1997). The first four subscales were also summed to provide a total difficulties score.

#### Cognitive Assessments

One subtest from the Phonological Assessment Battery was administered (PhAB; Frederickson et al., 1997). The Alliteration subtest measures the ability to isolate initial sounds of simple words. Raw scores from the PhAB Alliteration subtest were converted to standard scores (M = 100, SD = 15).

Four subtests from the Automated Working Memory Assessment were administered: Digit Recall, Backward Digit Recall, Dot Matrix, and Mr X (AWMA; Alloway, 2007). All are simple or complex memory tasks, with 6 trials at each span length. Digit Recall (testing verbal short-term memory) involves immediate serial recall of sequences of spoken digits; Backward Digit Recall follows the same procedure, except children are asked to recall items in reverse sequence. The Dot Matrix subtest (testing visuo-spatial short-term memory) requires children to recall the locations of a series of dots presented one at a time in a four-by-four matrix. The Mr X subtest (testing visuospatial working memory) involves recalling the location of a ball held by a cartoon character (‘Mr X’) for several successive displays involving different rotations of Mr X and ball positions. For all AWMA subtests, tasks automatically progress up one span level if a child produces four or more correct answers in a block. Subtests end following three or more incorrect responses. Trials correct were converted to age-standardised scores for each task (M = 100, SD = 15).

The Matrix Reasoning subtest of the Wechsler Abbreviated Scales of Intelligence II is used as an index of general reasoning and executive function (WASI; Wechsler, 2011). In this test, children are presented with incomplete matrices of images and asked to select an image that would suitably complete each matrix from a choice of four options. Trial numbers vary by age; children up to the age of 8 complete up to 24 matrices. Children aged 9 years and older complete a possible total of 30 matrices. The matrix reasoning test is finished when the child produces three consecutive incorrect responses. Trials correct were converted to T-scores (M = 10, SD = 10).

The Peabody Picture Vocabulary Test measures receptive vocabulary (PPVT; Dunn & Dunn, 2007). In this test, children were asked to select one image from four options according to what best corresponds to a stimulus word. In this test, a basal set is established after a child has completed all 12 items in set with one or no errors. Previous sets are presented in reverse-order until a basal set is successfully established. Children are presented with subsequent sets of increasing difficulty until a ‘ceiling’ set is established (defined by 8 or more errors out of twelve items). Raw scores were converted to standard scores (M = 100, SD = 15).

### Neuroimaging Data Acquisition

Magnetic Resonance Imaging (MRI) data were acquired at the MRC Cognition and Brain Sciences Unit, University of Cambridge. All scans were obtained on a Siemens 3T Prisma-Fit system (Siemens Healthcare, Erlangen, Germany) using a 32-channel head coil. T1-weighted volume scans were acquired using a whole-brain coverage 3D Magnetization Prepared Rapid Acquisition Gradient Echo (MP-RAGE) sequence acquired using 1-mm isometric image resolution (TR = 2.25s, TE = 2.98ms, flip angle = 9 degrees, 1×1×1mm). Diffusion scans were acquired using echo-planar diffusion-weighted images with a set of 60 non-collinear directions, using a weighting factor of b = 1000 s*mm−2, interleaved with a T2-weighted (b = 0) volume. Whole brain coverage was obtained with 60 contiguous axial slices and isometric image resolution of 2 mm. Echo time was 90ms and repetition time was 8400ms.

### Tractography and Connectome Construction

Pre-processing and reconstruction were performed using QSIPrep 0.14.2, which is based on Nipype 1.6.1 (nipype1; nipype2; Gorgolewski et al., 2011).

The T1-weighted (T1w) image was corrected for intensity non-uniformity (INU) using N4BiasFieldCorrection (n4; ANTs 2.3.1; Avants et al., 2014), and used as T1w-reference throughout the workflow. The T1w-reference was then skull-stripped using antsBrainExtraction.sh (ANTs 2.3.1), with OASIS as target template. Spatial normalization to the ICBM 152 Nonlinear Asymmetrical template version 2009c (MNI; Bowring et al., 2022) was performed through nonlinear registration with antsRegistration (ANTs 2.3.1), using brain-extracted versions of both T1w volume and template. Brain tissue segmentation of cerebrospinal fluid (CSF), white-matter (WM) and grey-matter (GM) was performed on the brain-extracted T1w using ‘FAST’ (FSL 6.0.3:b862cdd5; Jenkinson et al., 2011).

Any images with a b-value less than 100 s/mm^2 were treated as a b = 0 image. MP-PCA denoising as implemented in MRtrix3’s dwidenoise (dwidenoise1; Tournier et al., 2019) was applied with a 5-voxel window. After MP-PCA, B1 field inhomogeneity was corrected using dwibiascorrect from MRtrix3 with the N4 algorithm [n4]. After B1 bias correction, the mean intensity of the DWI series was adjusted so all the mean intensity of the b = 0 images matched across each separate DWI scanning sequence.

FSL (version 6.0.3:b862cdd5)’s Eddy was used for head motion correction and Eddy current correction. FSL’s Eddy was configured with a q-space smoothing factor of 10, a total of 5 iterations, and 1000 voxels to estimate hyperparameters. A linear first-level model and a linear second-level model were used to characterize Eddy current-related spatial distortion. Q-space coordinates were forcefully assigned to shells. Field offset was attempted to be separated from subject movement. Shells were aligned post-Eddy. Eddy’s outlier replacement was run [eddyrepol]. Data were grouped by slice, only including values from slices determined to contain at least 250 intracerebral voxels. Groups deviating by more than 4 standard deviations from the prediction had their data replaced with imputed values. Final interpolation was performed using the jac method.

Several confounding time-series were calculated based on the pre-processed DWI: framewise displacement (FD) using the implementation in Nipype (following the definitions by power_fd_dvars). The head-motion estimates calculated in the correction step were also placed within the corresponding confounds file. Slicewise cross-correlation was also calculated. The DWI time-series were resampled to ACPC, generating a pre-processed DWI run in ACPC space with 1mm isotropic voxels.

Diffusion orientation distribution functions were reconstructed using generalised q-sampling imaging (GQI; yeh2010gqi) with a ratio of mean diffusion distance of 1.250000.

Many internal operations of QSIPrep use Nilearn 0.8.0 (nilearn; Abraham et al., 2014) and Dipy. For more details of the pipeline, see QSIPrep’s documentation at https://qsiprep.readthedocs.io.

### Thresholding and Binarisation of Connectomes

Numerous different parcellation schemes can be used in the construction of connectomic models (e.g. DK-68, Desikan et al., 2016; AAL3, Rolls et al., 2020). In the current study, we used 100-node, Schaefer-parcellated streamline connectomes derived from DTI (Schaefer et al., 2018). We selected the Schaefer parcellation due to its functional correspondence to intrinsic connectivity networks in the brain. 100-node structural connectomes were binarised and then consistency-thresholded at 60%, in line with previous studies (e.g. de Reus & van den Heuvel, 2013). At a threshold of 39 streamlines (i.e. edges were required to have over 39 streamlines for us to assume an anatomical connection), a mean density of 9.18604% was calculated across connectomes. This approximate density value has previously been found to preserve the most informative, non-noisy edges in adjacency matrices, aiding the calculation of a range of graph-theoretic measures (van Wijk et al., 2010; Betzel et al., 2016; Akarca et al., 2021; Theis et al., 2021).

### Graph Theoretic Measures

Measures of brain networks can be represented in a number of ways, all of which capture different features of connectivity. In the current study, we used three graph theoretic measures at the nodal level in order to investigate the relationship between brain organisation and behaviour. These were nodal degree, clustering coefficient, and communicability (see Figure 2). To calculate these brain network properties across our binarised, consistency-thresholded 100-node streamline connectomes, we used Brain Connectivity Toolbox package in MATLAB (Rubinov & Sporns, 2010).

**Figure 1:**
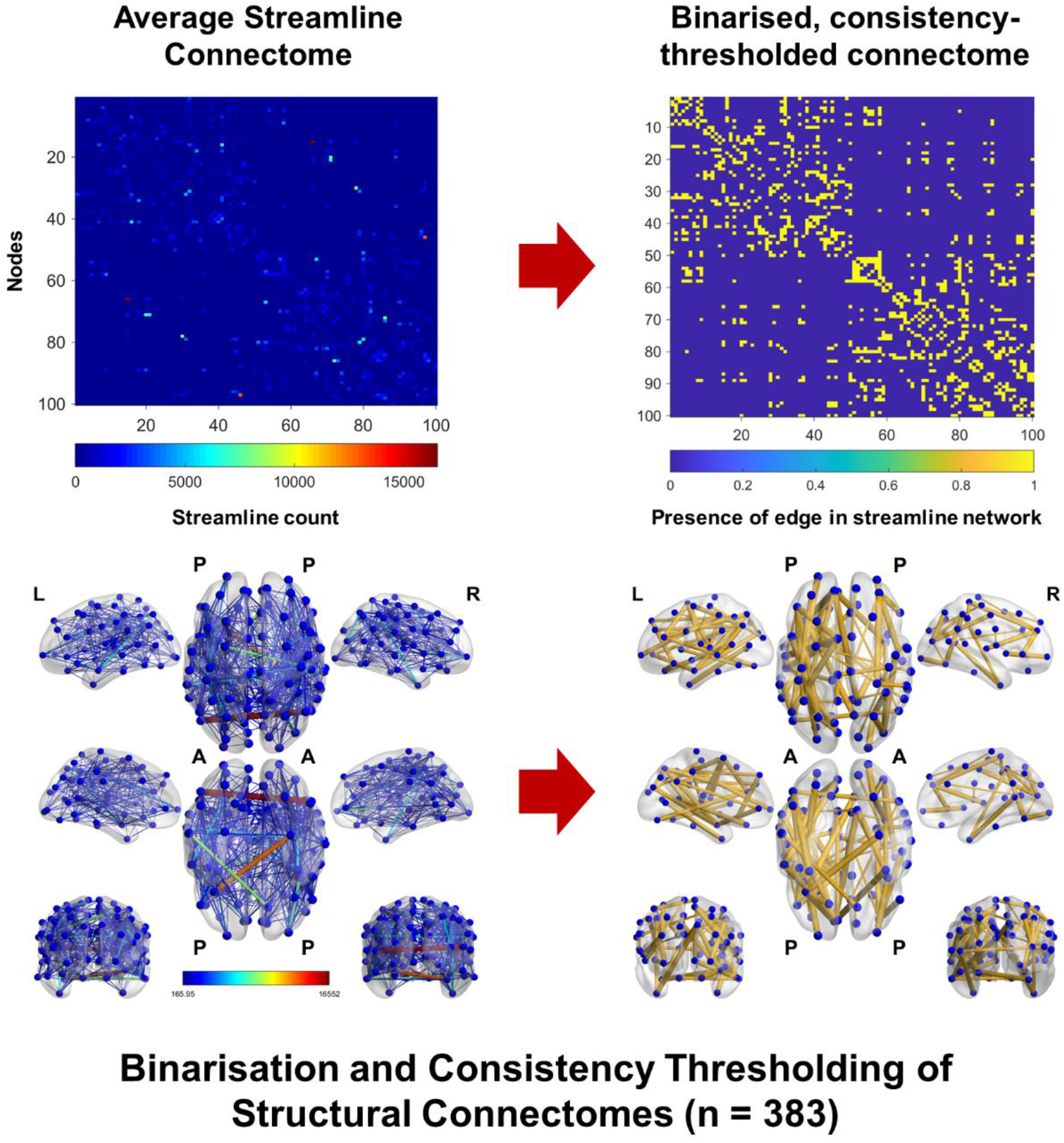
Adjacency matrix and brain network plot representing average streamline and binarised, consistency-thresholded connectomes across participants in our sample.

**Figure 2:**
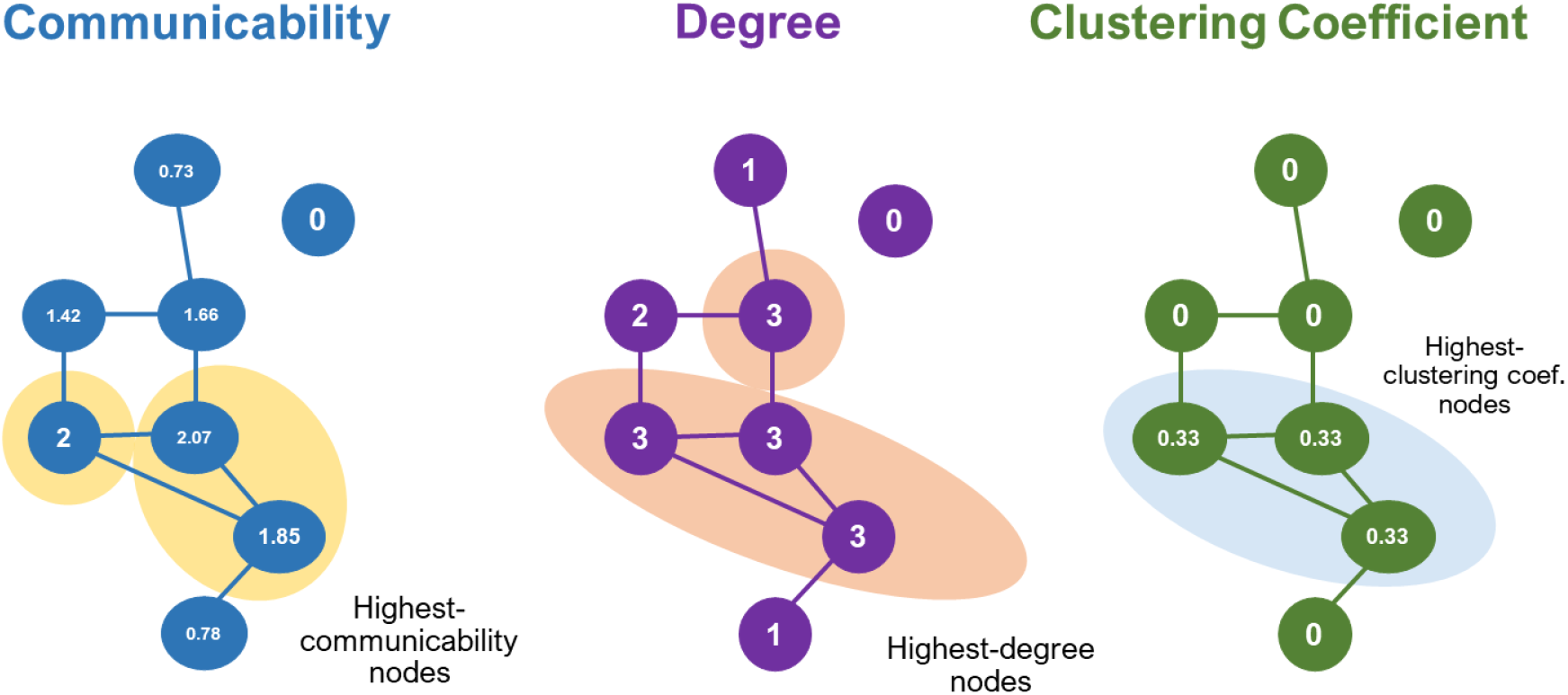
Diagram illustrating the properties of three nodal graph measures on an example network containing eight nodes and eight unweighted, undirected edges. Nodes with the highest communicability, degree, and clustering coefficient values are highlighted.

#### Nodal degree

The degree of an individual node is equal to the number of links connected to that node. In many real-world networks, including the brain, the distribution of degree values across nodes is heterogeneous—many nodes have a small number of links, and a few key nodes act as hubs of high connectivity that facilitate integration across the network (Fornito et al., 2016). Thus, the degree of a node—or brain region—serves as a basic measure of its integration within the broader brain network.

#### Local clustering coefficient

The clustering coefficient of a node quantifies how well-connected its neighbours are. Put simply, the clustering coefficient of a node represents the fraction of the node’s neighbours that are also neighbours of each other (Watts and Strogatz, 1998). Clustering coefficient values give insight into the community structure, or modularity, of a network. Since the brain is organised into structural and functional clusters of interconnected nodes, which strengthen into modules across the developmental timespan, the clustering coefficient is a valuable measure for gauging the contributions of individual nodes to the formation of modules (Meunier et al., 2010; Chen & Deem, 2015; Betzel et al., 2016; Betzel et al., 2017; Bassett & Sporns, 2017; Gallen & D’Esposito, 2019).

#### Communicability

The communicability of a node represents its direct and indirect connectedness to other brain regions. However, unlike betweenness centrality—which is calculated using shortest path lengths between nodes in a network—communicability is calculated using simulated random walks. Like betweenness centrality, communicability can be interpreted as a measure of a node’s ability to efficiently transmit information within a network (Funel, 2022). Communicability can also be thought of as the diffusion or propagation of a signal locally through a network. Since the brain develops in a way to minimise wiring costs and maximise information transfer, the efficiency with which brain regions are able to propagate a signal is a robust way of capturing this biological trade-off (Bullmore & Sporns, 2010; Power et al., 2010). Additionally, information transfer via short network paths and walks has been implicated in the development of psychiatric issues and cognitive difficulties, making it a useful measure for investigating the emergence of inattention and hyperactivity during childhood development (Seo et al., 2013; Makarov et al., 2018; Krukow et al., 2019; Silk et al., 2019).

## Analyses

### Exploratory Factor Analysis

We used Exploratory Factor Analysis (EFA) with a minimum-residual (‘minres’; also known as Iterated Principal Axis) estimation method to establish whether inattention and hyperactivity represent separate, or the same, underlying construct(s) in questionnaire data derived from the CALM sample. This was done using the factor_analyzer Python package (factor_analyzer version 0.4.0; Python version 3.9.7). No rotation was applied, as there was no specific aim to identify orthogonal underlying factors. Subscale scores from the Strengths and Difficulties Questionnaire (Hyperactivity subscale), Conners Questionnaire (Inattention and Hyperactivity/Impulsivity subscales), and BRIEF (Monitor and Working Memory subscales) were used. These questionnaires and their subscales have been found to have moderate-to-high internal consistency and reliability (Stone et al., 2015; Shaked et al., 2019; Zarrabi et al., 2015), and are regularly used in the clinical assessment of inattention and hyperactivity traits. While the BRIEF Monitor and Working Memory subscales do not directly measure hyperactivity and inattention, higher scores are significantly associated with having an ADHD diagnosis (Viola et al., 2018) and elevated inattention and hyperactivity (Jacobson et al., 2020). (Subscale scores were age-normalised and z-scored prior to the factor analysis.

### Partial Least Squares (PLS) Analyses

PLS, a data reduction technique similar to Principal Components Analysis, can adequately summarise the complex relationships between high-dimensional datasets (such as structural connectomes) and continuous variables (such as those representing variations in behaviour). PLS extracts orthogonal latent variables that maximally explain the covariance between predictor and response variables (Wold, 1975, 2004; Johnson et al., 2021). In our analysis, it was important to establish a link between variations in brain and those within behavioural data, such that significant variance in each could be explained by a common latent factor. PLS proved useful in achieving this, since it performs a dimensionality reduction that simultaneously considers patterns of variation in predictor variables and response variables, unlike PCA, which only uses principal components extracted from the predictor matrix as regressors on the response variable. The predictor variables for three separate PLS analyses were three node-level measures calculated using participants’ structural connectomes: nodal degree, clustering coefficient, and communicability. Participants’ inattention/hyperactivity factor scores were defined as the response variable (see Supplementary Figure 2 for a schematic representation of our PLS procedure).

One potential issue with PLS is the risk of overfitting. In order to control for this risk, we used recommendations made by Helmer et al. (2020) regarding the use of PLS to explore brain-behaviour relationships. We began by running 1000 permutations on participants’ nodal measures for each of our three PLS analyses. In our comparisons between empirical data and the probability distributions simulated using permutation, we calculated 95% confidence intervals using bootstrapped PLS outer weights (n = 1000, with replacement). This was done in order to test whether the bootstrapped confidence interval passed zero, and thus to establish which measures reliably load on the PLS components. For this bootstrapping we also applied a Procrustes rotation procedure to the outer weights. This was to account for sign flipping during the analysis (as previously recommended for PLS regressions; see Bastien, 2008).

### K-Means Clustering Analyses of Brain Data

Clustering is a data-driven approach that has previously been used to classify complex data, including both structural and functional whole-brain data (Rasero et al., 2019; Tokuda et al., 2021). We initially tested the strength of clustering solutions generated through hierarchical, k-means, and fuzzy clustering techniques. K-means clustering produced a solution with the highest silhouette value (the interpretation and implications of which we discuss further in this section). This technique carries additional benefits: k-means clustering scales to larger datasets and enables the identification of similarly-sized clusters, which allows for further statistical comparisons to be made between clusters (Jin et al., 2011). We performed k-means clustering on structural connectome data in order to establish whether our sample contains subgroups with differently-organised connectomes. This was done on the basis of three measures of brain network topology computed for nodes in binary, undirected connectomes: nodal degree, clustering coefficient, and communicability. These measures were z-score normalised across participants prior to the analysis.

We used metric multidimensional scaling, also known as principal coordinate analysis, to generate 2D representations of nodewise measures for each participant using the MDS package on MATLAB version 2019a. In generating these 2D representations, we used the robustcov function in MATLAB version 2019a to identify multivariate outliers, who were excluded from the analysis. Following this, we concatenated three 232 × 100 participant-by-node matrices, representing the three nodewise measures, to make a 232 × 300 participant-by-node matrix. We then calculated the Euclidean distances in these measures between participants, creating a 232×232 matrix. This enabled us to conduct our k-means clustering analysis on pairwise distances across each of the measures, maintaining dimensionality but projecting the data into 2D space. To determine the optimal number of clusters to extract in our k-means clustering analysis, we applied a silhouetting method, which assessed the separation between clusters across different solutions. Dalmaijer et al. (2022) calculated the statistical power of different k-means clustering solutions according to the number of clusters detected, sample sizes, and silhouette scores. To assess the strength of each of our potential clustering solutions, we used their criteria for statistical power, ensuring that the solution we included in further analyses detected clusters with high accuracy. Dalmaijer et al. (2022) determined that a k-means clustering solution with n ≥ 160 data points, and with a silhouette score Δ ≥ 5, has approximately 100% power to detect separation between clusters. A silhouette score threshold of Δ = 5 was therefore deemed suitable for establishing high statistical power and clustering accuracy, though we selected the clustering solution with the highest silhouette value. After determining that a two-cluster solution would be optimal according to this threshold (with a silhouette score of Δ = 6.808), we used two General Linear Models (GLMs) to explore how these two neural profiles differ along measures of behaviour and cognition. Head motion (measured by mean framewise displacement), age of scan, and age of test were included as regressors in the GLMs. To control for multiple comparisons, we applied a False Discovery Rate (FDR) correction with a 5% threshold to significant effects from these analyses.

Though our primary k-means clustering analysis was performed for the purpose of exploring neural heterogeneity in particularly inattentive and hyperactive children, we also ran a clustering analysis to investigate whether there are neural subtypes within our entire sample (n = 383). Supplementary Table 1 contains silhouette values for different *n*-cluster solutions across this broader sample.

**Table 1:** Table representing factor loadings for the questionnaires used in the factor analysis, assuming the existence of a single factor.

| **Questionnaire** | **Factor Loadings** | **Uniquenesses** |
| --- | --- | --- |
| **SDQ Hyperactivity** | 0.921 | 0.151 |
| **Conners Inattention** | 0.934 | 0.128 |
| **Conners Hyperactivity/Impulsivity** | 0.803 | 0.355 |
| **BRIEF Working Memory** | 0.896 | 0.198 |
| **BRIEF Monitor** | 0.845 | 0.286 |
| *Sum of Squares = 3.881* |  |  |

**Note.** Factor loadings represent weights and correlations between each variable and the factor. Uniqueness is the variance that is 'unique' to the variable and not shared with any of the other variables included in the factor analysis.

## Results

### Exploratory Factor Analysis

First, we used an EFA to determine whether inattention and hyperactivity represent one, or multiple, underlying factors, each explaining variance across multiple behavioural questionnaires. Data from five behavioural questionnaires measuring inattention and hyperactivity were used (see Methods section for further details). We found that a single Inattention/Hyperactivity factor explained 77.6% of the variance in questionnaire scores, indicating that these two features of behaviour tend to co-occur in the CALM sample (see Supplementary Figure 1 for scree plot representation of variance explained by *n* factors).

### PLS analysis does not show continuous neural dimensions that explain inattention and hyperactivity

We performed three separate PLS analyses on three node-level brain network measures: nodal degree, clustering coefficient, and communicability. In each of the three PLS analyses, the predictor matrix was therefore a 383 × 100 matrix (participant by node) where each element was the statistical property (degree, clustering coefficient, or communicability) and the response vector consisted of 383 × 1 factor loadings. Following our permutation procedure, the PLS analyses did not reveal any significant neural components that could explain variation along the behavioural continuum of inattention/hyperactivity in our sample (ppermuted > 0.2 for degree, clustering coefficient, and communicability). This demonstrates that although there might be a relationship between elements of brain organisation and inattention/hyperactivity, this cannot be reduced to linear underlying statistical components, at least with a sample of this size and composition.

### K-Means clustering reveals two groups with high inattention and hyperactivity

We restricted our next analysis to participants with clinically-elevated levels of inattention and hyperactivity, and then explored this variability in a different way. This new sample, which was characterised by elevated levels of inattention and hyperactivity (defined as surpassing a clinically-relevant threshold of a score over 60 on the Inattention and Hyperactivity/Impulsivity subscales of the Conners Questionnaire) left us with data from n = 232 children. These data were used in further analyses. In Supplementary Table 3, we compare the mean behavioural questionnaire and cognitive test scores of this group to those from the discarded sample (who had Conners Questionnaire scores <60; n = 151) and total sample (n = 383). Our clustering analysis tested whether individuals in this subsample, with high inattention and hyperactivity, are differentiated by three simple node-wise measures of their structural brain networks: degree, clustering coefficient, and communicability. We later explored which of these measures contributes the most to the clustering solution.

Following multidimensionality scaling across nodal measures, we calculated silhouette scores across a range of potential clustering solutions in order to find the optimal number of clusters to use in further analyses (see Supplementary Table 2 for alternative *n*-cluster solutions and corresponding silhouette values). The maximum silhouette score across these solutions was 0.6808. This clustering solution was represented by two similarly-sized clusters (n = 109 and n = 123).

### Differences in nodal degree, clustering coefficient, and communicability between the clusters

After identifying two groups of highly inattentive and hyperactive children using k-means clustering, we tested the extent to which the degree, clustering coefficient, and communicability of each node differentiated the groups. Put simply, having used these measures to identify the clusters, we wanted to test which nodewise measures were driving cluster membership. We compared each of these nodewise measures between clusters using general linear models, where participant age and MRI head motion (quantified across participants using mean framewise displacement) were included as control regressors. FDR post-hoc corrections were applied to account for multiple comparisons across 100 nodes for each of the graph measures (see Supplementary Figure 3 for a schematic of our GLM procedure, and Supplementary Figure 6 for frequency distributions of mean framewise displacement and age across participants).

### Higher Widespread Communicability, Clustering Coefficient, and Degree Values in Cluster 1

Since widespread nodal communicability appears to differentiate the clusters most strongly, we decided to investigate which of the clusters had the higher average communicability, clustering coefficient, and degree values across nodes. We conducted three paired-samples t-tests on averaged, z-score normalised communicability, clustering coefficient, and degree values. We found that Cluster 1 has, on average, higher nodal communicability (M = 0.6271, SD = 0.0851) than Cluster 2 (M = −0.6461, SD = 0.0623), t(198) = 120.6907, p = 2.3653 × 10^−187^, 95% C.I. [1.2524, 1.2940]. Additionally, Cluster 1 was found to have higher average nodal clustering coefficients (M = 0.0619, SD = 0.0930) than Cluster 2 (M = −0.0596, SD = 0.0903), t(198) = 9.3788, p = 1.5841 × 10^−17^, 95% CI [0.0960, 0.1471]. The final t-test revealed that Cluster 1 exhibits higher average nodal degree (M = 0.1317, SD = 0.1092) compared to Cluster 2 (M = −0.1412, SD = 0.1233), t(198) = 16.5693, p = 2.9489 × 10^−39^, 95% C.I. [0.2404, 0.3053].

### Cognitive differences, but not behavioural differences, characterise the neurotypes

In order to test whether our clustering solution generalises to behavioural and cognitive characteristics, we performed additional analyses comparing the clusters on nineteen different behavioural questionnaires and cognitive tests.

First, we used a Pearson’s Chi-Square Test to compare distributions of gender across groups. No significant differences in gender were found between the groups, X² (1, 232) = 0.521, p = 0.470. A paired-samples t-test was used to compare age (in months) between groups. Cluster 2 (M = 123.29, SD = 27.96) was found to be slightly older than Cluster 1 (M = 114.29, SD = 24.06), t(230) = −2.6111, p = 0.0096, 95% CI [−15.7898, −2.2084].

We used nineteen separate general linear models to test how behavioural and cognitive measures vary between the communicability-driven subtypes of those with clinically elevated inattention/hyperactivity. A GLM was performed for each behavioural and cognitive measure, and False Discovery Rate corrections were used to control for multiple comparisons across GLMs that included the same data type (for instance, across GLMs that included cognitive test scores). In all GLMs, age of scan, age of test, and mean framewise displacement (head motion) scores were included as control regressors (see Supplementary Figures 4 and 5 for schematic representations of our behavioural and cognitive GLM procedures).

The first set of GLMs, each of which included cluster membership as an independent variable and scores on one of twelve behavioural questionnaires as a dependent variable, tested how participants’ scores on behavioural questionnaires differed between the clusters. These questionnaires included the SDQ cumulative score, SDQ subscales (Emotion Regulation, Conduct, Hyperactivity, Peer Problems, and Prosocial), and Conners Questionnaire subscales (Inattention, Hyperactivity/Impulsivity, Learning Problems, Executive Function, and Aggression). Despite the fact that both clusters were already known to have clinically-elevated levels of inattention and hyperactivity, it was possible that there could be cluster-specific variations on inattention- and hyperactivity-related measures within the clinical range. However, no significant differences were found between the clusters on any of the questionnaire scales, t(227) < |1.9931|, p_corrected_ > 0.05.

The second set of GLMs, each of which included cluster membership as an independent variable and scores on one of seven cognitive tests as a dependent variable, tested cognitive test scores between the two clusters. Again, we corrected for multiple comparisons across tests using a False Discovery Rate correction. The cognitive tests included the WASI (Matrix Reasoning subtest), PPVT, PhAB (Alliteration subtest), and the AWMA (Dot Matrix, Backward Digit Recall, and Mr X subtests).

A significant difference was found in Weschler Abbreviated Scale of Intelligence (Matrix Reasoning subtest) scores between the clusters, t(227) = −2.3186, p_corrected_ = 0.0497. WASI scores in Cluster 1 (M = 45.68, SD = 10.03) were higher than those in Cluster 2 (M = 42.95, SD = 8.77). A significant difference was also found between the clusters’ scores on the Automated Working Memory Assessment (AWMA) Dot Matrix task, t(227) = −2.3665, p_corrected_ = 0.0497, and the AWMA Mr X task, t(227) = −2.4410, p_corrected_ = 0.0497. AWMA Dot Matrix scores in Cluster 1 (M = 93.48, SD = 17.46) were higher than those in Cluster 2 (M = 92.84, SD = 14.61), and that AWMA Mr X scores in Cluster 1 (M = 94.12, SD = 13.31) were higher than those in Cluster 2 (M = 91.52, SD = 12.36; see Figure 4). No significant differences were found between the clusters on the other cognitive tests, t(227) < |1.7690|, p_corrected_ > 0.05.

**Figure 3:**
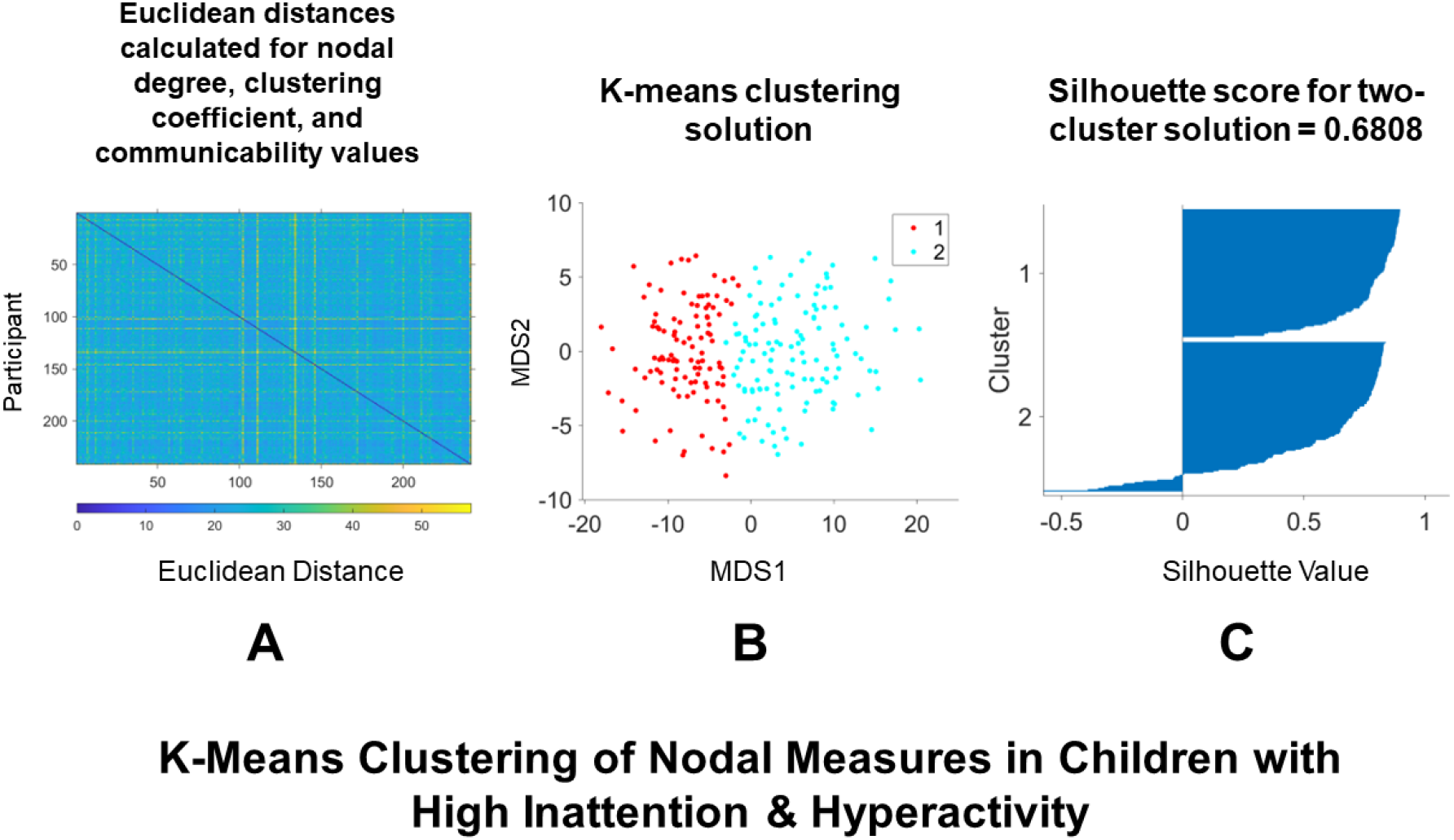
Diagram illustrating the steps we took in our k-means clustering analysis on graph theory measures derived from structural connectome data. We used multidimensionality scaling to generate a 2D Euclidean distance matrix between subjects (232 × 232), which was computed from nodal communicability, clustering coefficient, and degree values **(A)**. K-means clustering detected two groups, representing neural profiles based on these graph measures **(B)**. This two-cluster solution was selected due to its silhouette coefficient of 0.6808, indicating high degree of separation **(C)**.

**Figure 4:**
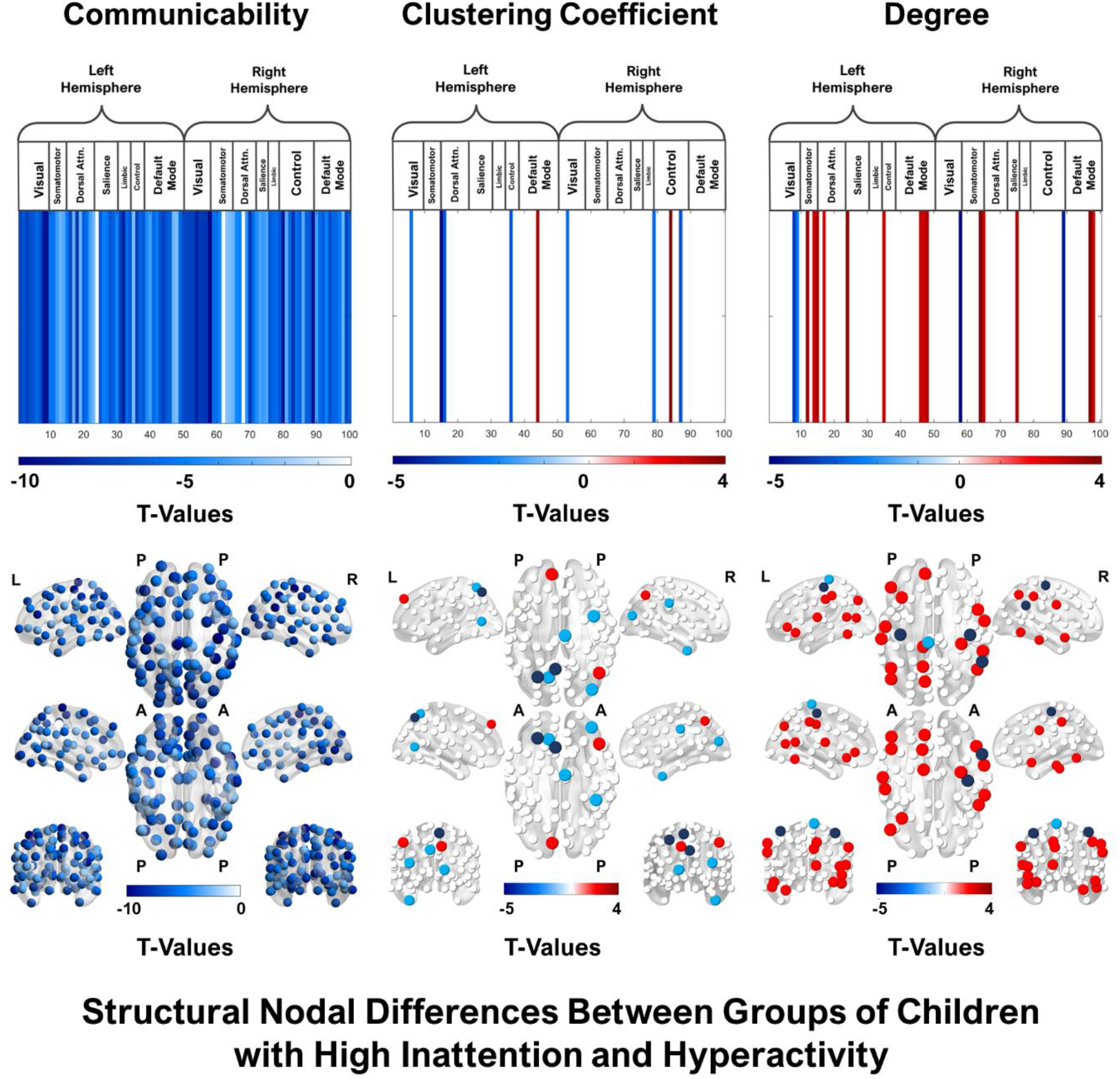
General linear models were used to compare nodewise measures between clusters of high inattention and hyperactivity. Differences were found in specific nodes across all three of the measures. The strongest and most widespread neural differences between the groups were reflected in nodal communicability values. A negative t-value indicates that the nodal measure is higher in Cluster 1 relative to Cluster 2.

**Figure 5:**
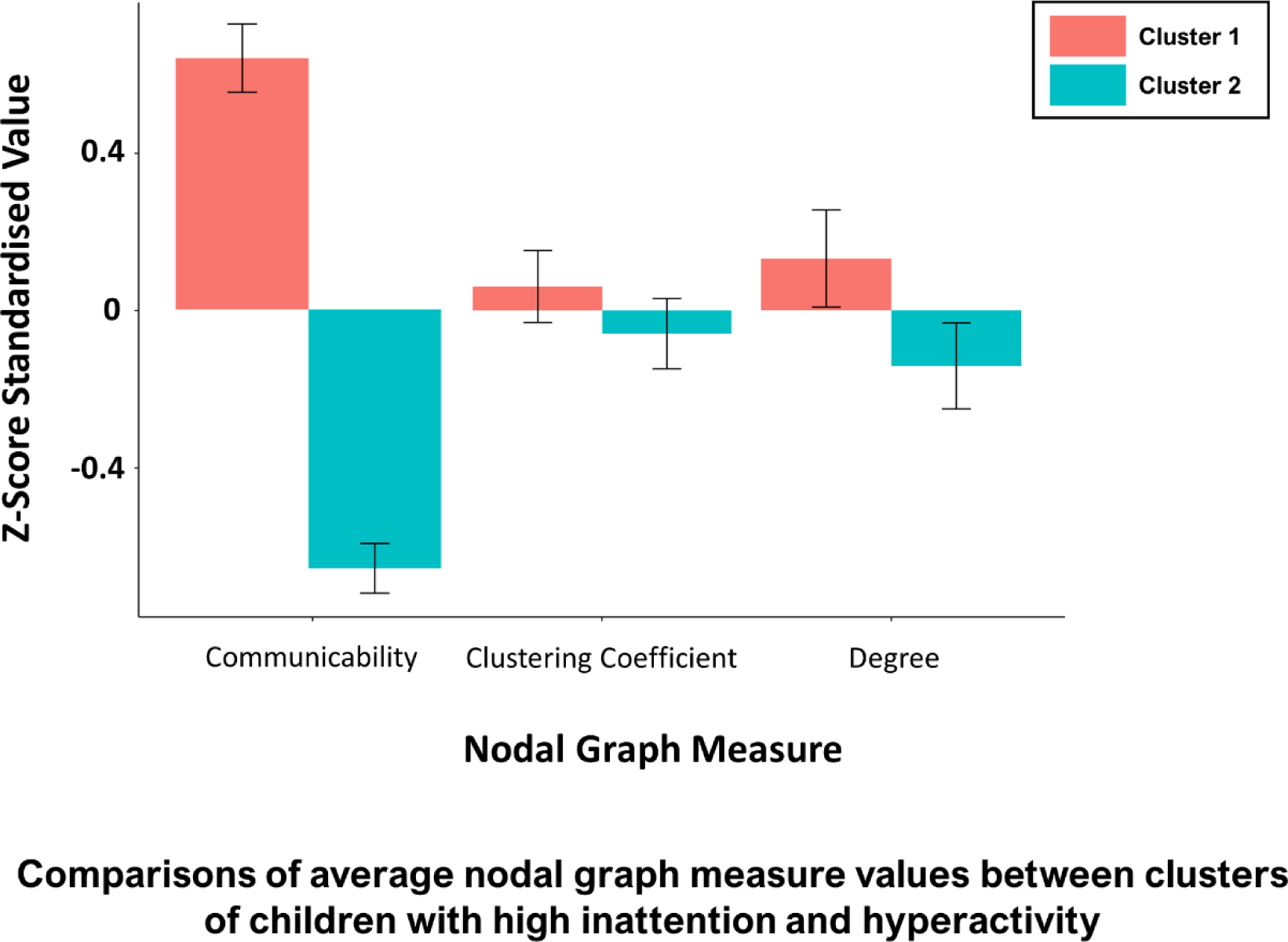
Paired-samples t-tests were used to compare global averages of nodewise measures between clusters of children with high inattention and hyperactivity. Children in Cluster 1 demonstrated higher nodal communicability, clustering coefficient, and degree values compared to Cluster 2. Error bars represent standard deviations of mean nodewise measure values.

**Figure 6:**
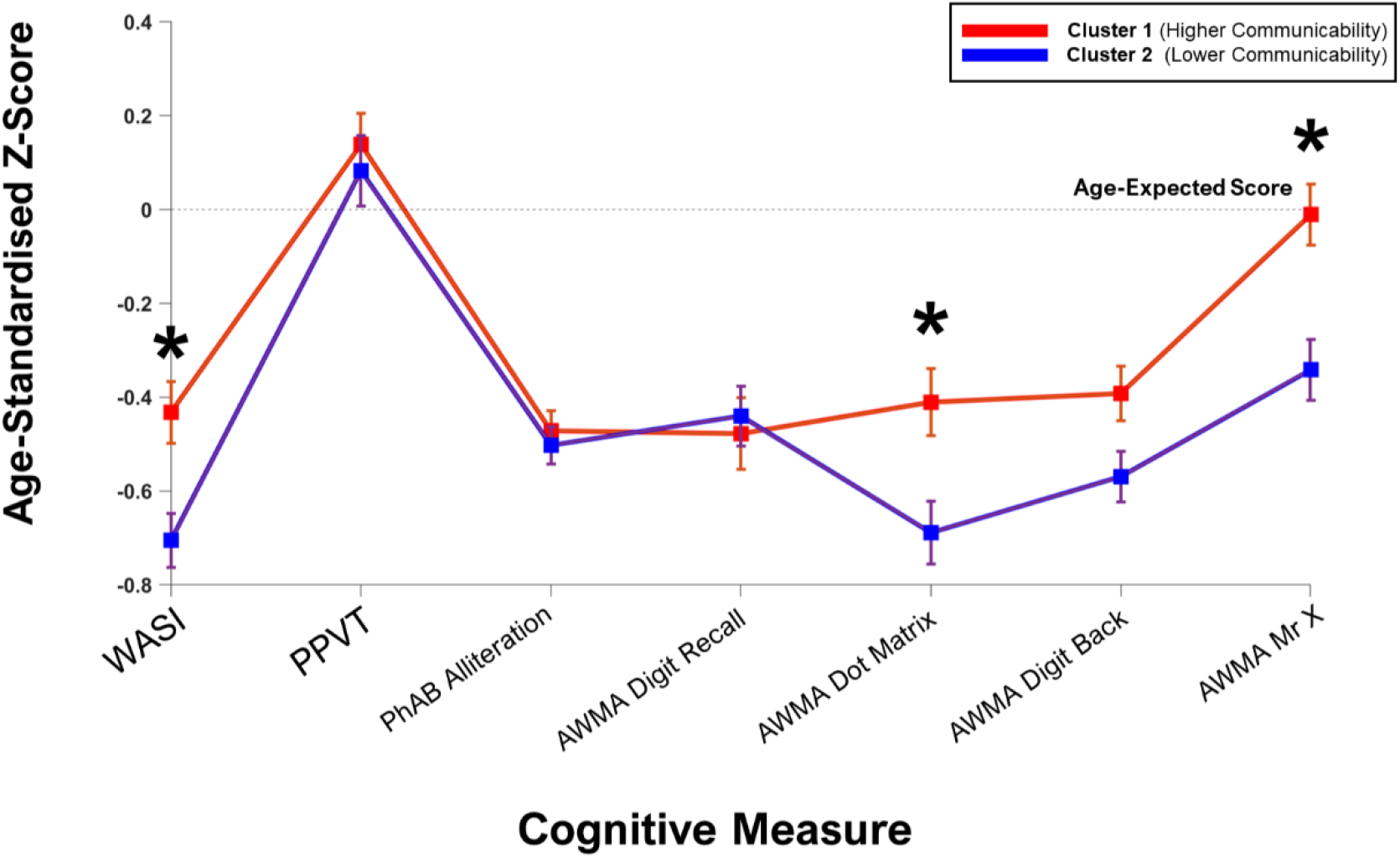
Age-standardised comparisons of cognitive differences between clusters based on cognitive test scores. Significant differences between clusters are labelled with an asterisk. **Note**. Error bars represent standard errors of mean cognitive test scores. Differences between the groups’ performance on cognitive tests were found on the Weschler Abbreviated Scale of Intelligence (WASI) and Automated Working Memory Assessment (AWMA; Dot Matrix and Mr X subtests).

## Discussion

We investigated whether, and how, differences in structural brain organisation mirrored a common transdiagnostic characteristic of children and young people with neurodevelopmental difficulties. Within our sample, inattention and hyperactivity were well-captured by a common latent factor, but this could not be well explained by linear connectomic differences. Across individuals with clinically-elevated inattention and hyperactivity, there was substantial heterogeneity in connectome organisation. We found that a two-cluster k-means clustering solution best captured this heterogeneity. These clusters of inattentive and hyperactive children were differentiated primarily by nodewise communicability, but also by clustering coefficient and degree. The two clusters had statistically-indistinguishable behavioural profiles—they were equally inattentive and hyperactive, and were comparable across various other developmentally-relevant domains, such as gender. However, these two groups did differ in their cognitive abilities, particularly on measures of executive function and visuospatial reasoning.

### A single dimension of inattention and hyperactivity

We first tested whether inattention and hyperactivity represent two separate behavioural constructs. We found that inattention and hyperactivity are highly related to one another within the CALM sample. One large inattention and hyperactivity factor can explain the majority of variance across subscales of the SDQ, Conners, and BRIEF questionnaires. This stands in contrast to accounts of inattention and hyperactivity that attribute these features of ADHD to separate behavioural ‘subtypes’ (August & Garfinkel, 1989). Indeed, there is increasing evidence of significant covariance between inattention and hyperactivity within ADHD, such that a general factor has better predictive power than two separate dimensions (Toplak et al., 2009; Sokolova et al., 2016). Additionally, Krakowski et al. (2020) found that variations in this latent inattention and hyperactivity factor do not differentiate the behavioural profiles of children with ADHD or autism, leading the authors of the study to suggest that this latent factor may also be more useful in the context of studying behaviour on a transdiagnostic basis.

It may be tempting to think that the demonstration of a common underlying factor for inattention and hyperactivity is somewhat at odds with the aims of transdiagnostic research. However, this does not logically follow. Determining the dimensional underpinnings of this characteristic is orthogonal to the question of the extent to which this factor extends across diagnostic boundaries, or co-occurs alongside other characteristics (Bathelt et al. 2018).

### Structural connectome properties did not linearly explain inattention and hyperactivity

Since inattention and hyperactivity can be explained by the same underlying construct in CALM, we chose to analyse brain network properties with this in mind, using these factor scores. We used three partial least squares regressions to test whether there are underlying components that represent co-variation between three node-level graph measures and inattention/hyperactivity factor scores. We did not find any significant components. This suggests that within the CALM sample, it may be difficult to explain the broad spectrum of inattention and hyperactivity with linear components relating to structural brain organisation. There are multiple possible explanations for this finding, one of them being sample size. With larger sample sizes, associations between these characteristics of structural connectivity and inattention/hyperactivity may be detectable, but the absence of effects here would suggest they are weak and/or highly variable. Another possibility is that differences in brain function, rather than brain structure, may be more valuable in defining linear variations in inattention and hyperactivity across a broad sample. A range of studies have investigated how measures of functional connectivity (FC) may differ in inattentive and hyperactive individuals compared to those who are ‘typically-developing’ (for review, see Castellanos & Aoki, 2016). Graph-theoretic measures are often used to inspect how these differences are manifested in patterns of FC, both across the whole brain and within, or between, specific networks or brain regions of interest. In CALM, Jones et al. (2022) found that hyperactivity/impulsivity was linearly associated with reduced segregation between higher order resting-state networks. Additionally, decreased global efficiency—an inverse measure of the topological distance between nodes in a network—has been found both in adults (Lin et al., 2014) and in children (Wang et al., 2009; Wang et al., 2020) who are inattentive and hyperactive. While the current study investigated structural brain organisation along a spectrum of inattention and hyperactivity, it is possible that further studies of FC in the CALM cohort will reveal differences in global and region-specific network measures that coincide with linear variations in behaviour.

A second explanation for the null results in the PLS analysis is that there are multiple different configurations of structural brain organisation that generate equally inattentive and hyperactive behavioural phenotypes. The current study lends evidence to the idea that inattention and hyperactivity do not result from one specific ‘type’ of brain, but that these features of behaviour are shared by multiple structural neurotypes. This is reflected in the concept of equifinality, which describes the means by which open systems reach similar observable end-states through different means (e.g. Simmons et al., 2019). The brain is an example of an open system—one which takes input from the outside world and engages in various embodied computational processes, resulting in observable cognitive and behavioural phenomena (Yufik et al., 2017). While there may be some consistencies in structural brain organisation across those with elevated inattention and hyperactivity (Gehricke et al., 2017; Bayard et al., 2020; Zhang-James et al., 2021), this does not discount the existence of heterogeneity in the neural structures and processes that could generate this behavioural phenotype. In a recent study of neuroanatomic heterogeneity in ADHD, Li et al. (2021) found that a range of neural subtypes existed within a large sample of individuals with ADHD in comparison to non-ADHD controls. The severity of inattention, hyperactivity, and impulsivity symptoms did not differ between these neural subtypes, suggesting that there is no stable, one-to-one mapping between brain types and complex behavioural types. Additionally, a recent meta-analysis conducted by Cortese et al. (2021) found no consistent differences in functional connectivity across individuals with ADHD compared to controls, which may be accounted for by heterogeneity in study participants. The current study supports these findings in the context of a transdiagnostic framework, which enabled us to go beyond procedures that draw discrete distinctions between those with and without a diagnosis of ADHD.

### Two clusters of brain organisation within inattention and hyperactivity

Because we did not find a linear component that explained variations in brain organisation along a broad spectrum of inattention and hyperactivity, we decided to explore possible nonlinearities between these behavioural traits and attributes of brain organisation. To investigate neurological diversity among children with particularly high levels of inattention and hyperactivity, we ran a k-means clustering analysis on the three nodal measures from the structural connectomes of children whose Conners Questionnaire t-scores exceeded the clinical threshold of 60 on both of the relevant Conners Questionnaire subscales (Inattention and Hyperactivity/Impulsivity). The cleanest clustering solution produced two clusters of structural brain organisation (n = 109 and 123). Admittedly, one can see that the clustering algorithm is essentially carving a continuous distribution in half. Nonetheless, the silhouette score indicates that there is indeed a good separation between the two clusters. This suggests that these two groups can be distinguished on the basis of high or low MDS scores but that the boundary may not be well-defined.

The presence of neural heterogeneity, which can be characterised (albeit coarsely) as two clusters, is a demonstration that multiple different neural profiles coincide with the expression of clinically significant levels of inattention and hyperactivity. Our comparisons of nodal degree, clustering coefficient, and communicabilitiy across the two clusters revealed that all three of these measures pick up on nodewise differences between the groups. However, communicability shows the greatest nodewise differences, suggesting that they are best differentiated by this feature of structural brain network organisation.

Communicability is a representation of theoretical information flow through a network, based on the number of short-path connections that link a single node to other nodes (Benzi et al., 2013). The differentiation of children using this measure has interesting implications within the context of brain development and neural-behavioural equifinality (Crofts & Higham, 2009; Andreotti et al., 2014). The first implication hinges on the widespread relevance of communicability across whole-brain networks in differentiating subgroups of inattentive and hyperactive children. The brain structure of one group of children is characterised by almost universally higher-communicability nodes than the other. However, both of these groups exhibit scores on the Conners Questionnaire that exceed the clinically-relevant threshold. Different neural subtypes can exist among children who express the same degree of behavioural inattention and hyperactivity, demonstrating the principle of neural-behavioural equifinality. Brain network topologies that facilitate more energetically-efficient information transfer support the development of more robust frontoparietal networks, which in turn are thought give rise to differences in executive function (Cui et al., 2020). Additionally, polysynaptic network communication models, captured by communicability, have been shown to be predictive of human functional and behavioural characteristics, over and above models that do not capture this mode of connectivity (Seguin & Zalesky, 2020; Seguin et al., 2022). Our findings demonstrate that the structural architecture of whole-brain networks may play a role in directing cognitive and behavioural processes, and could provide new insights into the anatomical basis for developmental diversity. It is possible that communicability, alongside other measures of brain network topology, relates to a set of closely-associated cognitive capacities underpinned by a broader developmental process. Since communicability measures the ability of a node to propagate a signal through a network, increases in this neural measure may be expected to coincide with the development of more efficient forms of cognitive processing. Because differences in cognitive ability are widely seen across neurodevelopmental conditions, structural brain network communicability—and the biological mechanisms by which it changes over time—may provide a window into how these differences emerge.

In the current study, we show that group-based differences in structural brain organisation may have consequences for cognitive development. Although both clusters have statistically-indistinguishable behavioural profiles, they differ significantly on three cognitive assessments measuring visuospatial reasoning, spatial short-term memory, and spatial working memory. Children in Cluster 1 scored higher than children in Cluster 2 on all three measures. Higher widespread nodal communicability, clustering coefficient, and degree values also characterises children from Cluster 1. This could suggest a link between global patterns of structural brain organisation, measured across nodes, and cognitive measures of executive function in children with elevated inattention and hyperactivity. ADHD has previously been associated with differences in inter- and intra-network connectivity across multiple brain networks, with additional evidence pointing towards global reductions in network-specific integration and segregation (Cao et al., 2016; Qian et al., 2019; Hilger & Fiebach, 2019; Griffiths et al., 2021; Jones et al., 2022). Across infancy and childhood, structural brain networks serve as an anatomical backbone to the development of functional modules (Giedd & Rapoport, 2010). These networks undergo a process of gradual refinement, characterised by changes in modular integration and segregation according to a rich-club topological structure that is well-established by the time of birth (Ball et al., 2014; Kim & Min, 2020). Over time, structural brain modules, or *hubs*, develop short-path connections to other modules, enabling efficient long-range communication across brain networks (Park & Friston, 2016). As such, brain development is characterised by an optimisation in the balance between the global integration and local segregation of information transfer, which is revealed by features of structural brain network topologies. A number of studies have observed characteristic associations between intelligence and node-specific measures of within- and between-module connectivity, particularly in frontal and parietal brain regions (Gallen et al., 2016; Hilger et al., 2017; Chaddock & Heyman, 2020). Developmental differences in the emergence and interconnectivity of these structural modules may affect the type, rate, and degree of cognitive and behavioural change across childhood. In the case of children with elevated inattention and hyperactivity, widespread decreases in nodal degree and clustering, which has implications for connectivity within and between structural modules, may coincide with lower cognitive performance, as observed in Cluster 2. Additionally, multiple studies have identified nodal communicability as a relevant measure in the association between brain organisation and cognitive ability (Gilson et al., 2020; Lella & Vessio, 2021). This could reflect a trend of reduced communication across networks in the brain, indicated by the formation of fewer short paths between brain areas. However, the current study did not examine the directionality of the causal mechanisms underpinning differences in nodal communicability. Future longitudinal research might be able to reveal the temporal sequencing of the relationship between cognitive performance and communicability, shedding light on how these related measures change throughout development.

As discussed previously, the concurrent presentation of inattention and hyperactivity in our sample enabled us to explore these behavioural measures through one underlying dimension. The cognitive differences between children in Cluster 1 and Cluster 2 indicate that these neurotypes also reflect two phenotypic subtypes of high inattention and hyperactivity. Since we applied k-means clustering to participants’ connectomes based on a clinical threshold, all of these children could arguably qualify for a clinical diagnosis of ADHD. However, they were not distinguished by inattention *or* hyperactivity—subtypes which exist in clinical definitions of ADHD—but by their cognitive ability, which also coincided with differences in their structural brain network topologies. This has potential implications for the treatment of ADHD, particularly in the provision of subtype-specific accommodations in educational settings. Previously, Reiersen and Todorov (2013) found that current diagnostic criteria may fail to identify a subgroup of individuals with both inattention and hyperactivity who may have higher support needs than other individuals with an ADHD diagnosis. We argue that these heightened support needs may emerge from not only ADHD-specific symptoms, like inattention and hyperactivity, but the fact that a sizeable subgroup of children with ADHD also experience cognitive difficulties that act as enhanced barriers to learning in school. It is crucial that future research investigates the specific attributes and educational needs of children with differing cognitive difficulties, as this can give a more precise direction to educational policymaking and resource allocation.

### Limitations

One potential limitation of the current study was its sample size. An initial dataset of n = 383 may have hindered the detection of linear components in brain organisation that could explain variance in inattention and hyperactivity. This was further reduced to n = 232 in analyses of participants with clinically elevated inattention and hyperactivity. Having more data available for our analyses may have allowed us to identify more specific subgroups in our clustering analysis—perhaps with different, or more subtle, variations in cognition and behaviour. In the future, data from larger developmental cohorts should be incorporated into studies of brain network differences in the domain of heightened inattention and hyperactivity.

Additionally, our use of k-means clustering ensured that clusters were defined along a hard boundary, such that groups of a similar size were maintained in our clustering solution. Other methods, such as gaussian mixture modelling, have previously been used to identify the extent to which individual datapoints hold ‘responsibility’ within their respective clusters (McLachlan, 2004; Reynolds, 2009; Yang et al., 2012). This allows datapoints to be sorted along a gradient of contribution to a clustering solution, rather than strictly allocated to a particular cluster. In the future, we recommend that these methods be applied to ascertain the validity and reliability of clustering solutions—particularly those which are more ambiguous in their statistical power.

Following the identification of n = 232 children with elevated inattention and hyperactivity in our larger sample, we were left with an additional subset of n = 151 children whose Conners Questionnaire scores did not exceed 60. While considerations had been made about the possibility of using this subset as a ‘control’ group, and including these children into our comparisons between inattentive and hyperactive clusters, we made the decision to only retain participants who surpassed the clinical threshold. While the Conners Questionnaire defines a clinical threshold of >60 for the purpose of identifying children with ADHD-like characteristics, it does not define a range of scores for children who are ‘typically-developing.’ In the future, it may be possible to define hallmark features of ‘typical’ behavioural and cognitive development. However, this taxonomical project would also be challenging, since these features fall along their own severity spectrums, and significant heterogeneity exists between individuals (see Astle & Fletcher-Watson, 2020, for review). In Supplementary Table 3, we summarise the behavioural and cognitive characteristics of children across our whole sample, compared to those in the inattentive/hyperactive and Conners Questionnaire<60 subsamples.

The fact that the inattentive and hyperactive clusters were not found to differ on any behavioural measures could be attributable to the fact that functional connectivity is typically better-associated with behaviour than structural connectivity (Jia et al., 2014; Fjell et al., 2017). In addition, structural brain connectivity has been linked robustly to cognitive ability (Ponsoda et al., 2017; Babaeeghazvini et al., 2021; Amunts et al., 2022). This coupling of structure-cognition and function-behaviour relationships could explain why cognitive differences, rather than behavioural differences, were found to exist between two different profiles of structural brain connectivity. Future studies should strive to investigate whether there are subgroups of inattentive and hyperactive children that are differentiable by their functional connectivity profiles, and whether additional contrasts on other behavioural measures can be observed between these groups.

## Conclusion

We found that properties of structural brain networks differ among children with high levels of inattention and hyperactivity. Additionally, we were able to see how two neurotypes with the same behavioural presentation differ on measures of cognitive ability, which may represent a relevant dimension of assessment when characterising the needs of children with inattention and hyperactivity. A next step for research in this field might be to investigate whether, and how, properties of functional connectivity could differentiate subgroups of highly inattentive and hyperactive individuals. Studies that clarify these relationships will provide further insight into potential links between attributes of structural and functional brain organisation, as well as cognition and behaviour, in children with elevated inattention and hyperactivity. The current study represents a methodological and theoretical step towards the use of transdiagnostic approaches in the context of studying the development of the human connectome.

## Acknowledgements

The authors were supported by the Medical Research Council program grant MC-A0606-5PQ41, and D.E.A. and D.A. were supported by an Opportunity Award from the James S. McDonnell Foundation. We would like to thank all members of the CALM Team for their help with recruitment, data collection, and data management, as well as all of the children and parents for their participation in the study. The CALM Team includes lead investigators Duncan Astle, Kate Baker, Rogier Kievit and Tom Manly. Data collection was assisted by a team of researchers and PhD students including Danyal Akarca, Joe Bathelt, Marc Bennett, Madalena Bettencourt, Giacomo Bignardi, Sarah Bishop, Erica Bottacin, Lara Bridge, Diandra Brkic, Annie Bryant, Sally Butterfield, Elizabeth Byrne, Gemma Crickmore, Edwin Dalmaijer, Fánchea Daly, Tina Emery, Laura Forde, Grace Franckel, Delia Furhmann, Andrew Gadie, Sara Gharooni, Jacalyn Guy, Erin Hawkins, Agnieszka Jaroslawska, Sara Joeghan, Amy Johnson, Jonathan Jones, Silvana Mareva, Elise Ng-Cordell, Sinead O’Brien, Cliodhna O’Leary, Joseph Rennie, Ivan Simpson-Kent, Roma Siugzdaite, Tess Smith, Stephani Uh, Maria Vedechkina, Francesca Woolgar, Natalia Zdorovtsova, and Mengya Zhang. The authors wish to thank the many professionals working in children’s services in the South-East and East of England for their support, and to the children and their families for giving up their time to visit the clinic. We would also like to thank the radiographers who support the excellent paediatric scanning at the MRC CBSU.

## Conflict of Interest Statement

The authors declare no conflicts of interest.

## Notes

### Competing Interest Statement

The authors have declared no competing interest.

## References

Abraham, A., Pedregosa, F., Eickenberg, M., Gervais, P., Mueller, A., Kossaifi, J., Gramfort, A., Thirion, B., & Varoquaux, G. (2014). Machine learning for neuroimaging with scikit-learn. Frontiers in Neuroinformatics, 8(FEB), 14. https://doi.org/10.3389/FNINF.2014.00014/BIBTEX

Akarca, D., Vértes, P. E., Bullmore, E. T., Baker, K., Gathercole, S. E., Holmes, J., Kievit, R. A., Manly, T., Bathelt, J., Bennett, M., Bignardi, G., Bishop, S., Bottacin, E., Bridge, L., Brkic, D., Bryant, A., Butterfield, S., Byrne, E. M., Crickmore, G., … Astle, D. E. (2021). A generative network model of neurodevelopmental diversity in structural brain organization. Nature Communications 2021 12:1, 12(1), 1–18. https://doi.org/10.1038/s41467-021-24430-z

Alloway TP. Automated working memory assessment. London, UK: Pearson; 2007.

Alloway, T., Gathercole, S. E., Kirkwood, H., & Elliott, J. (2011). Evaluating the validity of the Automated Working Memory Assessment. https://Doi.Org/10.1080/01443410802243828, 28(7), 725–734. https://doi.org/10.1080/01443410802243828

Amunts, K., Defelipe, J., Pennartz, C., Destexhe, A., Migliore, M., Ryvlin, P., Furber, S., Knoll, A., Bitsch, L., Bjaalie, J. G., Ioannidis, Y., Lippert, T., Sanchez-Vives, M. V., Goebel, R., & Jirsa, V. (2022). Linking Brain Structure, Activity, and Cognitive Function through Computation. ENeuro, 9(2). https://doi.org/10.1523/ENEURO.0316-21.2022

Andreotti, J., Jann, K., Melie-Garcia, L., Giezendanner, S., Dierks, T., & Federspiel, A. (2014). Repeatability Analysis of Global and Local Metrics of Brain Structural Networks. https://Home.Liebertpub.Com/Brain, 4(3), 203–220. https://doi.org/10.1089/BRAIN.2013.0202

Astle, D. E., Bathelt, J., & Holmes, J. (2019). Remapping the cognitive and neural profiles of children who struggle at school. Developmental Science, 22(1), e12747. https://doi.org/10.1111/DESC.12747

Astle, D. E., Holmes, J., Kievit, R., & Gathercole, S. E. (2022). Annual Research Review: The transdiagnostic revolution in neurodevelopmental disorders. Journal of Child Psychology and Psychiatry, 63(4), 397–417. https://doi.org/10.1111/JCPP.13481

Astle, D. E., & Fletcher-Watson, S. (2020). Beyond the Core-Deficit Hypothesis in Developmental Disorders. Current Directions in Psychological Science, 29(5), 431–437. https://doi.org/10.1177/0963721420925518

August, G., & Garfinkel, B. (1989). Behavioral and Cognitive Subtypes of ADHD. Journal of the American Academy of Child and Adolescent Psychiatry, 28(5), 739–748. https://doi.org/10.1097/00004583-198909000-00016

Avants, B. B., Tustison, N., & Johnson, H. (2014). Advanced Normalization Tools (ANTS) Release 2.x. https://brianavants.wordpress.com/2012/04/13/updated-ants-compile-instructions-april-12-2012/

Avena-Koenigsberger, A., Misic, B., & Sporns, O. (2017). Communication dynamics in complex brain networks. Nature Reviews. Neuroscience, 19(1), 17–33. https://doi.org/10.1038/NRN.2017.149

Babaeeghazvini, P., Rueda-Delgado, L. M., Gooijers, J., Swinnen, S. P., & Daffertshofer, A. (2021). Brain Structural and Functional Connectivity: A Review of Combined Works of Diffusion Magnetic Resonance Imaging and Electro-Encephalography. Frontiers in Human Neuroscience, 15, 585. https://doi.org/10.3389/FNHUM.2021.721206/BIBTEX

Ball, G., Aljabar, P., Zebari, S., Tusor, N., Arichi, T., Merchant, N., Robinson, E. C., Ogundipe, E., Rueckert, D., Edwards, A. D., & Counsell, S. J. (2014). Rich-club organization of the newborn human brain. Proceedings of the National Academy of Sciences of the United States of America, 111(20), 7456–7461. https://doi.org/10.1073/PNAS.1324118111/-/DCSUPPLEMENTAL

Bassett, D. S., & Sporns, O. (2017). Network neuroscience. Nature Neuroscience, 20(3), 353–364. https://doi.org/10.1038/NN.4502

Bastien, P. (2008). Deviance residuals based PLS regression for censored data in high dimensional setting. Chemometrics and Intelligent Laboratory Systems, 91(1), 78–86. https://doi.org/10.1016/J.CHEMOLAB.2007.09.009

Bathelt, J., Gathercole, S. E., Butterfield, S., & Astle, D. E. (2018). Children’s academic attainment is linked to the global organization of the white matter connectome. Developmental Science, 21(5), e12662. https://doi.org/10.1111/DESC.12662

Bayard, F., Nymberg Thunell, C., Abé, C., Almeida, R., Banaschewski, T., Barker, G., Bokde, A. L. W., Bromberg, U., Büchel, C., Quinlan, E. B., Desrivières, S., Flor, H., Frouin, V., Garavan, H., Gowland, P., Heinz, A., Ittermann, B., Martinot, J. L., Martinot, M. L. P., … Petrovic, P. (2020). Distinct brain structure and behavior related to ADHD and conduct disorder traits. Molecular Psychiatry, 25(11), 3020. https://doi.org/10.1038/S41380-018-0202-6

Benzi, M., Klymko, C., & Estrada, E. (2013). Total communicability as a centrality measure. Journal of Complex Networks, 1(2), 124–149. https://doi.org/10.1093/COMNET/CNT007

Betzel, R. F., Avena-Koenigsberger, A., Goñi, J., He, Y., de Reus, M. A., Griffa, A., Vértes, P. E., Mišic, B., Thiran, J. P., Hagmann, P., van den Heuvel, M., Zuo, X. N., Bullmore, E. T., & Sporns, O. (2016). Generative models of the human connectome. NeuroImage, 124(Pt A), 1054–1064. https://doi.org/10.1016/J.NEUROIMAGE.2015.09.041

Betzel, R. F., & Bassett, D. S. (2017). Multi-scale brain networks. NeuroImage, 160, 73–83. https://doi.org/10.1016/J.NEUROIMAGE.2016.11.006

Betzel, R. F., Cutts, S. A., Greenwell, S., Faskowitz, J., & Sporns, O. (2022). Individualized event structure drives individual differences in whole-brain functional connectivity. NeuroImage, 252. https://doi.org/10.1016/J.NEUROIMAGE.2022.118993

Bos, D. J., Merchán-Naranjo, J., Martínez, K., Pina-Camacho, L., Balsa, I., Boada, L., Schnack, H., Oranje, B., Desco, M., Arango, C., Parellada, M., Durston, S., & Janssen, J. (2015). Reduced Gyrification Is Related to Reduced Interhemispheric Connectivity in Autism Spectrum Disorders. Journal of the American Academy of Child and Adolescent Psychiatry, 54(8), 668–676. https://doi.org/10.1016/J.JAAC.2015.05.011

Bowring, A., Nichols, T. E., & Maumet, C. (2022). Isolating the sources of pipeline-variability in group-level task-fMRI results. Human Brain Mapping, 43(3), 1112–1128. https://doi.org/10.1002/HBM.25713

Bullmore, E., & Sporns, O. (2012). The economy of brain network organization. Nature Reviews. Neuroscience, 13(5), 336–349. https://doi.org/10.1038/NRN3214

Buckholtz, J. W., & Meyer-Lindenberg, A. (2012). Psychopathology and the human connectome: toward a transdiagnostic model of risk for mental illness. Neuron, 74(6), 990–1004. https://doi.org/10.1016/J.NEURON.2012.06.002

Campbell, J. M., & Dommestrup, A. K. (2010). Peabody Picture Vocabulary Test. The Corsini Encyclopedia of Psychology, 1–1. https://doi.org/10.1002/9780470479216.CORPSY0649

Cao, M., Huang, H., Peng, Y., Dong, Q., & He, Y. (2016). Toward Developmental Connectomics of the Human Brain. Frontiers in Neuroanatomy, 10, 25. https://doi.org/10.3389/FNANA.2016.00025/BIBTEX

Castellanos, F. X., & Aoki, Y. (2016). Intrinsic Functional Connectivity in Attention-Deficit/Hyperactivity Disorder: A Science in Development. Biological Psychiatry. Cognitive Neuroscience and Neuroimaging, 1(3), 253–261. https://doi.org/10.1016/J.BPSC.2016.03.004

Chaddock-Heyman, L., Weng, T. B., Kienzler, C., Weisshappel, R., Drollette, E. S., Raine, L. B., Westfall, D. R., Kao, S. C., Baniqued, P., Castelli, D. M., Hillman, C. H., & Kramer, A. F. (2020). Brain Network Modularity Predicts Improvements in Cognitive and Scholastic Performance in Children Involved in a Physical Activity Intervention. Frontiers in Human Neuroscience, 14. https://doi.org/10.3389/FNHUM.2020.00346

Chen, M., & Deem, M. W. (2015). Development of modularity in the neural activity of children’s brains. Physical Biology, 12(1), 016009. https://doi.org/10.1088/1478-3975/12/1/016009

Coghill, D., & Sonuga-Barke, E. J. S. (2012). Annual research review: categories versus dimensions in the classification and conceptualisation of child and adolescent mental disorders--implications of recent empirical study. Journal of Child Psychology and Psychiatry, and Allied Disciplines, 53(5), 469–489. https://doi.org/10.1111/J.1469-7610.2011.02511.X

Connaughton, M., Whelan, R., O’Hanlon, E., & McGrath, J. (2022). White matter microstructure in children and adolescents with ADHD. NeuroImage : Clinical, 33, 102957. https://doi.org/10.1016/J.NICL.2022.102957

Conners, C. K., Pitkanen, J., & Rzepa, S. R. (2011). Conners 3rd Edition (Conners 3; Conners 2008). Encyclopedia of Clinical Neuropsychology, 675–678. https://doi.org/10.1007/978-0-387-79948-3_1534

Cortese, S., Aoki, Y. Y., Itahashi, T., Castellanos, F. X., & Eickhoff, S. B. (2021). Systematic Review and Meta-analysis: Resting-State Functional Magnetic Resonance Imaging Studies of Attention-Deficit/Hyperactivity Disorder. Journal of the American Academy of Child & Adolescent Psychiatry, 60(1), 61–75. https://doi.org/10.1016/J.JAAC.2020.08.014

Costa, L. D. F., Rodrigues, F. A., Travieso, G., & Boas, P. R. V. (2007). Characterization of complex networks: A survey of measurements. Advances in Physics, 56(1), 167–242. https://doi.org/10.1080/00018730601170527

Crofts, J. J., & Higham, D. J. (2009). A weighted communicability measure applied to complex brain networks. Journal of the Royal Society Interface, 6(33), 411. https://doi.org/10.1098/RSIF.2008.0484

Cui, Z., Stiso, J., Baum, G. L., Kim, J. Z., Roalf, D. R., Betzel, R. F., Gu, S., Lu, Z., Xia, C. H., He, X., Ciric, R., Oathes, D. J., Moore, T. M., Shinohara, R. T., Ruparel, K., Davatzikos, C., Pasqualetti, F., Gur, R. E., Gur, R. C., … Satterthwaite, T. D. (2020). Optimization of energy state transition trajectory supports the development of executive function during youth. ELife, 9, 1–60. https://doi.org/10.7554/ELIFE.53060

Cuthbert, B. N. (2014). The RDoC framework: facilitating transition from ICD/DSM to dimensional approaches that integrate neuroscience and psychopathology. World Psychiatry, 13(1), 28. https://doi.org/10.1002/WPS.20087

Dalgleish, T., Black, M., Johnston, D., & Bevan, A. (2020). Transdiagnostic Approaches to Mental Health Problems: Current Status and Future Directions. Journal of Consulting and Clinical Psychology, 88(3), 179. https://doi.org/10.1037/CCP0000482

Dalmaijer, E. S., Nord, C. L., & Astle, D. E. (2022). Statistical power for cluster analysis. BMC Bioinformatics, 23(205). https://doi.org/10.1186/s12859-022-04675-1

de Reus, M. A., & van den Heuvel, M. P. (2013). The parcellation-based connectome: limitations and extensions. NeuroImage, 80, 397–404. https://doi.org/10.1016/J.NEUROIMAGE.2013.03.053

Dellapiazza, F., Michelon, C., Vernhet, C., Muratori, F., Blanc, N., Picot, M. C., Baghdadli, A., Baghdadli, A., Chabaux, C., Chatel, C., Cohen, D., Damville, E., Geoffray, M. M., Gicquel, L., Jardri, R., Maffre, T., Novo, A., Odoyer, R., Oreve, M. J., … Vespirini, S. (2021). Sensory processing related to attention in children with ASD, ADHD, or typical development: results from the ELENA cohort. European Child & Adolescent Psychiatry, 30(2), 283–291. https://doi.org/10.1007/S00787-020-01516-5

Desikan, R. S., Ségonne, F., Fischl, B., Quinn, B. T., Dickerson, B. C., Blacker, D., Buckner, R. L., Dale, A. M., Maguire, R. P., Hyman, B. T., Albert, M. S., & Killiany, R. J. (2006). An automated labeling system for subdividing the human cerebral cortex on MRI scans into gyral based regions of interest. NeuroImage, 31, 968–980. https://doi.org/10.1016/j.neuroimage.2006.01.021

Dunn, L.M., & Dunn, D. M. (2007). Peabody Picture Vocabulary Test. Pearson Education: Minneapolis, USA.

Jeste, S. S. & Geschwind, D. H. (2014). Disentangling the heterogeneity of autism spectrum disorder through genetic findings. Nat. Rev. Neurol, 10, 74–81. https://doi.org/10.1038/nrneurol.2013.278

Fjell, A. M., Sneve, M. H., Grydeland, H., Storsve, A. B., Amlien, I. K., Yendiki, A., & Walhovd, K. B. (2017). Relationship between structural and functional connectivity change across the adult lifespan: A longitudinal investigation. Human Brain Mapping, 38(1), 561. https://doi.org/10.1002/HBM.23403

Forest, D. (2014). Neuroconstructivism: A Developmental Turn in Cognitive Neuroscience?. In: Wolfe, C.T. (eds) Brain Theory. Palgrave Macmillan, London. https://doi.org/10.1057/9780230369580_5

Fornito, A., Zalesky, A., & Bullmore, E. T. (2016). Fundamentals of Brain Network Analysis. In Fundamentals of Brain Network Analysis. Elsevier Inc. https://doi.org/10.1016/C2012-0-06036-X

Foss-Feig, J. H., Fontaine, N. de la, & Tsatsanis, K. (2016). BRIEF (Behavior Rating Inventory of Executive Functions). Encyclopedia of Autism Spectrum Disorders, 1–5. https://doi.org/10.1007/978-1-4614-6435-8_102048-1

Frederickson, N. (1997). Phonological assessment battery (PhAB) : manual and test materials. NFER-Nelson.

Funel, A. (2022). A method to compute the communicability of nodes through causal paths in temporal networks. Physica A: Statistical Mechanics and Its Applications, 593, 126965. https://doi.org/10.1016/J.PHYSA.2022.126965

Fusar-Poli, P., Solmi, M., Brondino, N., Davies, C., Chae, C., Politi, P., Borgwardt, S., Lawrie, S. M., Parnas, J., & McGuire, P. (2019). Transdiagnostic psychiatry: a systematic review. World Psychiatry : Official Journal of the World Psychiatric Association (WPA), 18(2), 192–207. https://doi.org/10.1002/WPS.20631

Gallen, C. L., Baniqued, P. L., Chapman, S. B., Aslan, S., Keebler, M., Didehbani, N., & D’Esposito, M. (2016). Modular Brain Network Organization Predicts Response to Cognitive Training in Older Adults. PloS One, 11(12). https://doi.org/10.1371/JOURNAL.PONE.0169015

Gallen, C. L., & D’Esposito, M. (2019). Brain Modularity: A Biomarker of Intervention-related Plasticity. Trends in Cognitive Sciences, 23(4), 293–304. https://doi.org/10.1016/J.TICS.2019.01.014

Gehricke, J. G., Kruggel, F., Thampipop, T., Alejo, S. D., Tatos, E., Fallon, J., & Muftuler, L. T. (2017). The brain anatomy of attention-deficit/hyperactivity disorder in young adults – a magnetic resonance imaging study. PLoS ONE, 12(4). https://doi.org/10.1371/JOURNAL.PONE.0175433

Gilmore, J. H., Knickmeyer, R. C., & Gao, W. (2018). Imaging structural and functional brain development in early childhood. Nature Reviews. Neuroscience, 19(3), 123. https://doi.org/10.1038/NRN.2018.1

Giedd, J. N., & Rapoport, J. L. (2010). Structural MRI of Pediatric Brain Development: What Have We Learned and Where Are We Going? Neuron, 67(5), 728–734. https://doi.org/10.1016/J.NEURON.2010.08.040

Gilson, M., Zamora-López, G., Pallarés, V., Adhikari, M. H., Senden, M., Campo, A. T., Mantini, D., Corbetta, M., Deco, G., & Insabato, A. (2020). Model-based whole-brain effective connectivity to study distributed cognition in health and disease. Network Neuroscience, 4(2), 338. https://doi.org/10.1162/NETN_A_00117

Gioia, G. A., Isquith, P. K., Guy, S. C., & Kenworthy, L. (2000). Behaviour Rating Inventory of Executive Function - Ages 5–18 (BRIEF) Psychological Assessment Resources: Florida, USA.

Goodman, R. (1997). The Strengths and Difficulties Questionnaire: a research note. Journal of Child Psychology and Psychiatry, and Allied Disciplines, 38(5), 581–586. https://doi.org/10.1111/J.1469-7610.1997.TB01545.X

Gorgolewski, K., Burns, C. D., Madison, C., Clark, D., Halchenko, Y. O., Waskom, M. L., & Ghosh, S. S. (2011). Nipype: a flexible, lightweight and extensible neuroimaging data processing framework in python. Frontiers in Neuroinformatics, 5, 13. https://doi.org/10.3389/fninf.2011.00013

Griffiths, D. L., Farrell, L. J., Waters, A. M., & White, S. W. (2017). Clinical correlates of obsessive compulsive disorder and comorbid autism spectrum disorder in youth. Journal of Obsessive-Compulsive and Related Disorders, 14, 90–98. https://doi.org/10.1016/J.JOCRD.2017.06.006

Griffiths, K. R., Braund, T. A., Kohn, M. R., Clarke, S., Williams, L. M., & Korgaonkar, M. S. (2021). Structural brain network topology underpinning ADHD and response to methylphenidate treatment. Translational Psychiatry, 11(1), 150. https://doi.org/10.1038/S41398-021-01278-X

Gui, A., Mason, L., Gliga, T., Hendry, A., Begum Ali, J., Pasco, G., Shephard, E., Curtis, C., Charman, T., Johnson, M. H., Meaburn, E., & Jones, E. J. H. (2020). Look duration at the face as a developmental endophenotype: Elucidating pathways to autism and ADHD. Development and Psychopathology, 32(4), 1303–1322. https://doi.org/10.1017/S0954579420000930

Hayiou-Thomas, M. E., Smith-Woolley, E., & Dale, P. S. (2021). Breadth versus depth: Cumulative risk model and continuous measure prediction of poor language and reading outcomes at 12. Developmental Science, 24(1), e12998. https://doi.org/10.1111/DESC.12998

Helmer, M., Warrington, S., Mohammadi-Nejad, A.-R., Ji, J. L., Howell, A., Rosand, B., Anticevic, A., Sotiropoulos, S. N., & Murray, J. D. (2020). On stability of Canonical Correlation Analysis and Partial Least Squares with application to brain-behavior associations. BioRxiv. https://doi.org/10.1101/2020.08.25.265546

Hilger, K., Ekman, M., Fiebach, C. J., & Basten, U. (2017). Intelligence is associated with the modular structure of intrinsic brain networks. Scientific Reports 2017 7:1, 7(1), 1–12. https://doi.org/10.1038/s41598-017-15795-7

Hilger, K., & Fiebach, C. J. (2019). ADHD symptoms are associated with the modular structure of intrinsic brain networks in a representative sample of healthy adults. Network Neuroscience, 3(2), 567. https://doi.org/10.1162/NETN_A_00083

Holmes, J., Bryant, A., & Gathercole, S. E. (2019). Protocol for a transdiagnostic study of children with problems of attention, learning and memory (CALM) 17 Psychology and Cognitive Sciences 1701 Psychology 17 Psychology and Cognitive Sciences 1702 Cognitive Sciences 11 Medical and Health Sciences 1117 Public Health and Health Services. BMC Pediatrics, 19(1), 1–11. https://doi.org/10.1186/S12887-018-1385-3/FIGURES/1

Jacob, S., Wolff, J. J., Steinbach, M. S., Doyle, C. B., Kumar, V., & Elison, J. T. (2019). Neurodevelopmental heterogeneity and computational approaches for understanding autism. Translational Psychiatry 2019 9:1, 9(1), 1–12. https://doi.org/10.1038/s41398-019-0390-0

Jenkinson, M., Beckmann, C. F., Behrens, T. E. J., Woolrich, M. W., & Smith, S. M. (2012). FSL. NeuroImage, 62(2), 782–790. https://doi.org/10.1016/J.NEUROIMAGE.2011.09.015

Jia, H., Hu, X., & Deshpande, G. (2014). Behavioral Relevance of the Dynamics of the Functional Brain Connectome. Brain Connectivity, 4(9), 741. https://doi.org/10.1089/BRAIN.2014.0300

Jin X., Han J. (2011) K-Means Clustering. In: Sammut C., Webb G.I. (eds) Encyclopedia of Machine Learning. Springer, Boston, MA. https://doi.org/10.1007/978-0-387-30164-8_425

Johnson, A., Bathelt, J., Akarca, D., & Astle, D. E. (2021). Far and wide: Associations between childhood socio-economic status and brain connectomics. Developmental Cognitive Neuroscience, 48, 100888. https://doi.org/10.1016/J.DCN.2020.100888

Jones, J. S., Astle, D. E., Baker, K., Gathercole, S., Holmes, J., Kievit, R., Manly, T., Akarca, D., Bathelt, J., Bennett, M., Bettencourt, M., Bignardi, G., Bishop, S., Bottacin, E., Bridge, L., Brkic, D., Bryant, A., Butterfield, S., Byrne, E., … Zhang, M. (2022). Segregation and integration of the functional connectome in neurodevelopmentally “at risk” children. Developmental Science, 25(3). https://doi.org/10.1111/DESC.13209

Jones, J. S., Team, T. C., Leyland-Craggs, A., & Astle, D. E. (2022). Testing the Triple Network Model of Psychopathology in a Transdiagnostic Neurodevelopmental Cohort. MedRxiv, 2022.05.05.22274709. https://doi.org/10.1101/2022.05.05.22274709

Joshi, G., Wozniak, J., Petty, C., Martelon, M. K., Fried, R., Bolfek, A., Kotte, A., Stevens, J., Furtak, S. L., Bourgeois, M., Caruso, J., Caron, A., & Biederman, J. (2013). Psychiatric comorbidity and functioning in a clinically referred population of adults with autism spectrum disorders: a comparative study. Journal of Autism and Developmental Disorders, 43(6), 1314–1325. https://doi.org/10.1007/S10803-012-1679-5

Kern, J. K., Geier, D. A., King, P. G., Sykes, L. K., Mehta, J. A., & Geier, M. R. (2015). Shared Brain Connectivity Issues, Symptoms, and Comorbidities in Autism Spectrum Disorder, Attention Deficit/Hyperactivity Disorder, and Tourette Syndrome. Brain Connectivity, 5(6), 321–335. https://doi.org/10.1089/BRAIN.2014.0324

Kim, D. J., & Min, B. K. (2020). Rich-club in the brain’s macrostructure: Insights from graph theoretical analysis. Computational and Structural Biotechnology Journal, 18, 1761. https://doi.org/10.1016/J.CSBJ.2020.06.039

Kofler, M. J., Irwin, L. N., Soto, E. F., Groves, N. B., Harmon, S. L., & Sarver, D. E. (2019). Executive Functioning Heterogeneity in Pediatric ADHD. Journal of Abnormal Child Psychology, 47(2), 273–286. https://doi.org/10.1007/S10802-018-0438-2

Konrad, A., Dielentheis, T. F., El Masri, D., Bayerl, M., Fehr, C., Gesierich, T., Vucurevic, G., Stoeter, P., & Winterer, G. (2010). Disturbed structural connectivity is related to inattention and impulsivity in adult attention deficit hyperactivity disorder. The European Journal of Neuroscience, 31(5), 912–919. https://doi.org/10.1111/J.1460-9568.2010.07110.X

Krakowski, A. D., Cost, K. T., Anagnostou, E., Lai, M. C., Crosbie, J., Schachar, R., Georgiades, S., Duku, E., & Szatmari, P. (2020). Inattention and hyperactive/impulsive component scores do not differentiate between autism spectrum disorder and attention-deficit/hyperactivity disorder in a clinical sample. Molecular Autism, 11(1), 1–13. https://doi.org/10.1186/S13229-020-00338-1/FIGURES/2

Krukow, P., Jonak, K., Karpiński, R., & Karakuła-Juchnowicz, H. (2019). Abnormalities in hubs location and nodes centrality predict cognitive slowing and increased performance variability in first-episode schizophrenia patients. Scientific Reports 2019 9:1, 9(1), 1–13. https://doi.org/10.1038/s41598-019-46111-0

Lau-Zhu, A., Fritz, A., & McLoughlin, G. (2019). Overlaps and distinctions between attention deficit/hyperactivity disorder and autism spectrum disorder in young adulthood: Systematic review and guiding framework for EEG-imaging research. Neuroscience and Biobehavioral Reviews, 96, 93–115. https://doi.org/10.1016/J.NEUBIOREV.2018.10.009

Lella, E., & Estrada, E. (2020). Communicability distance reveals hidden patterns of Alzheimer’s disease. Network Neuroscience, 4(4), 1007–1029. https://doi.org/10.1162/NETN_A_00143

Li, T., van Rooij, D., Roth Mota, N., Buitelaar, J. K., Hoogman, M., Arias Vasquez, A., Franke, B., Ambrosino, S., Banaschewski, T., Bandeira, C. E., Bau, C. H. D., Baumeister, S., Baur-Streubel, R., Bellgrove, M. A., Biederman, J., Bralten, J., Bramati, I. E., Brandeis, D., Berm, S., … Schwarz, L. (2021). Characterizing neuroanatomic heterogeneity in people with and without ADHD based on subcortical brain volumes. Journal of Child Psychology and Psychiatry, 62(9), 1140–1149. https://doi.org/10.1111/JCPP.13384

Lin, P., Sun, J., Yu, G., Wu, Y., Yang, Y., Liang, M., & Liu, X. (2014). Global and local brain network reorganization in attention-deficit/hyperactivity disorder. Brain Imaging and Behavior, 8(4), 558–569. https://doi.org/10.1007/S11682-013-9279-3/TABLES/4

Makarov, V. V, Zhuravlev, M. O., Runnova, A. E., Protasov, P., Maksimenko, V. A., Frolov, N. S., Pisarchik, A. N., & Hramov, A. E. (2018). Betweenness centrality in multiplex brain network during mental task evaluation. PHYSICAL REVIEW E, 98, 62413. https://doi.org/10.1103/PhysRevE.98.062413

Mareva, S., & Holmes, J. (2019). Transdiagnostic associations across communication, cognitive, and behavioural problems in a developmentally at-risk population: A network approach. BMC Pediatrics, 19(1), 1–12. https://doi.org/10.1186/S12887-019-1818-7/FIGURES/3

MATLAB. (2018). 9.7.0.1190202 (R2019b). Natick, Massachusetts: The MathWorks Inc.

McClain, M. B., Harris, B., Schwartz, S. E., Benallie, K. J., Golson, M. E., & Benney, C. M. (2019). Brief Report: Development and Validation of the Autism Spectrum Knowledge Scale General Population Version: Preliminary Analyses. Journal of Autism and Developmental Disorders, 49(7), 3007–3015. https://doi.org/10.1007/S10803-019-04019-8

McLachlan, G. J., & Chang, S. U. (2004). Mixture modelling for cluster analysis. Statistical Methods in Medical Research, 13(5), 347–361. https://doi.org/10.1191/0962280204sm372ra

Meunier, D., Lambiotte, R., & Bullmore, E. T. (2010). Modular and hierarchically modular organization of brain networks. Frontiers in Neuroscience, 4(DEC), 200. https://doi.org/10.3389/FNINS.2010.00200/BIBTEX

Mizuno, Y., Kagitani-Shimono, K., Jung, M., Makita, K., Takiguchi, S., Fujisawa, T. X., Tachibana, M., Nakanishi, M., Mohri, I., Taniike, M., & Tomoda, A. (2019). Structural brain abnormalities in children and adolescents with comorbid autism spectrum disorder and attention-deficit/hyperactivity disorder. Translational Psychiatry 2019 9:1, 9(1), 1–7. https://doi.org/10.1038/s41398-019-0679-z

Newby, J. M., McKinnon, A., Kuyken, W., Gilbody, S., & Dalgleish, T. (2015). Systematic review and meta-analysis of transdiagnostic psychological treatments for anxiety and depressive disorders in adulthood. Clinical Psychology Review, 40, 91–110. https://doi.org/10.1016/J.CPR.2015.06.002

Nigg, J. (2013). Attention-deficit/hyperactivity disorder and adverse health outcomes. Clinical Psychology Review, 33(2), 215–228. https://doi.org/10.1016/J.CPR.2012.11.005

Park, H. J., & Friston, K. (2013). Structural and functional brain networks: From connections to cognition. Science, 342(6158). https://doi.org/10.1126/SCIENCE.1238411

Pereira, A. M., Campos, B. M., Coan, A. C., Pegoraro, L. F., de Rezende, T. J. R., Obeso, I., Dalgalarrondo, P., da Costa, J. C., Dreher, J. C., & Cendes, F. (2018). Differences in cortical structure and functional MRI connectivity in high functioning autism. Frontiers in Neurology, 9(JUL), 539. https://doi.org/10.3389/FNEUR.2018.00539/FULL

Ponsoda, V., Martínez, K., Pineda-Pardo, J. A., Abad, F. J., Olea, J., Román, F. J., Barbey, A. K., & Colom, R. (2017). Structural brain connectivity and cognitive ability differences: A multivariate distance matrix regression analysis. Human Brain Mapping, 38(2), 803–816. https://doi.org/10.1002/HBM.23419

Power, J. D., Cohen, A. L., Nelson, S. M., Wig, G. S., Barnes, K. A., Church, J. A., Vogel, A. C., Laumann, T. O., Miezin, F. M., Schlaggar, B. L., & Petersen, S. E. (2011). Functional network organization of the human brain. Neuron, 72(4), 665. https://doi.org/10.1016/J.NEURON.2011.09.006

Qian, X., Castellanos, F. X., Uddin, L. Q., Loo, B. R. Y., Liu, S., Koh, H. L., Poh, X. W. W., Fung, D., Guan, C., Lee, T. S., Lim, C. G., & Zhou, J. (2019). Large-scale brain functional network topology disruptions underlie symptom heterogeneity in children with attention-deficit/hyperactivity disorder. NeuroImage: Clinical, 21, 101600. https://doi.org/10.1016/J.NICL.2018.11.010

Rasero, J., Diez, I., Cortes, J. M., Marinazzo, D., & Stramaglia, S. (2019). Connectome sorting by consensus clustering increases separability in group neuroimaging studies. Network Neuroscience, 3(2), 325. https://doi.org/10.1162/NETN_A_00074

Reiersen, A. M., & Todd, R. D. (2008). Co-occurrence of ADHD and autism spectrum disorders: phenomenology and treatment. Expert Review of Neurotherapeutics, 8(4), 657–669. https://doi.org/10.1586/14737175.8.4.657

Reiersen, A. M., & Todorov, A. A. (2013). Exploration of ADHD Subtype Definitions and Co-Occurring Psychopathology in a Missouri Population-Based Large Sibship Sample. Scandinavian Journal of Child and Adolescent Psychiatry and Psychology, 1(1), 3–13. https://doi.org/10.21307/SJCAPP-2013-002

Reynolds, D. (2009). Gaussian Mixture Models. Encyclopedia of Biometrics, 659–663. https://doi.org/10.1007/978-0-387-73003-5_196

Rolls, E. T., Huang, C. C., Lin, C. P., Feng, J., & Joliot, M. (2020). Automated anatomical labelling atlas 3. NeuroImage, 206, 116189. https://doi.org/10.1016/J.NEUROIMAGE.2019.116189

Roth, R. M., & Saykin, A. J. (2004). Executive dysfunction in attention-deficit/hyperactivity disorder: cognitive and neuroimaging findings. The Psychiatric Clinics of North America, 27(1), 83–96. https://doi.org/10.1016/S0193-953X(03)00112-6

Rubinov, M., & Sporns, O. (2010). Complex network measures of brain connectivity: Uses and interpretations. NeuroImage, 52(3), 1059–1069. https://doi.org/10.1016/J.NEUROIMAGE.2009.10.003

Schaefer, A., Kong, R., Gordon, E. M., Laumann, T. O., Zuo, X. N., Holmes, A. J., Eickhoff, S. B., & Yeo, B. (2018). Local-Global Parcellation of the Human Cerebral Cortex from Intrinsic Functional Connectivity MRI. Cerebral cortex (New York, N.Y. : 1991), 28(9), 3095–3114. https://doi.org/10.1093/cercor/

Seguin, C., Tian, Y., & Zalesky, A. (2020). Network communication models improve the behavioral and functional predictive utility of the human structural connectome. Network Neuroscience (Cambridge, Mass.), 4(4), 980–1006. https://doi.org/10.1162/NETN_A_00161

Seo, E. H., Lee, D. Y., Lee, J. M., Park, J. S., Sohn, B. K., Lee, D. S., Choe, Y. M., & Woo, J. I. (2013). Whole-brain Functional Networks in Cognitively Normal, Mild Cognitive Impairment, and Alzheimer’s Disease. PLOS ONE, 8(1), e53922. https://doi.org/10.1371/JOURNAL.PONE.0053922

Shaked, D., Faulkner, L. M. D., Tolle, K., Wendell, C. R., Waldstein, S. R., & Spencer, R. J. (2019). Reliability and validity of the Conners’ Continuous Performance Test. https://Doi.Org/10.1080/23279095.2019.1570199, 27(5), 478–487. https://doi.org/10.1080/23279095.2019.1570199

Shephard, E., Bedford, R., Milosavljevic, B., Gliga, T., Jones, E. J. H., Pickles, A., Johnson, M. H., Charman, T., Baron-Cohen, S., Bolton, P., Chandler, S., Elsabbagh, M., Fernandes, J., Garwood, H., Hudry, K., Pasco, G., Tucker, L., & Volein, A. (2019). Early developmental pathways to childhood symptoms of attention-deficit hyperactivity disorder, anxiety and autism spectrum disorder. Journal of Child Psychology and Psychiatry, and Allied Disciplines, 60(9), 963–974. https://doi.org/10.1111/JCPP.12947

Silk, T. J., Malpas, C. B., Beare, R., Efron, D., Anderson, V., Hazell, P., Jongeling, B., Nicholson, J. M., & Sciberras, E. (2019). A network analysis approach to ADHD symptoms: More than the sum of its parts. PLOS ONE, 14(1), e0211053. https://doi.org/10.1371/JOURNAL.PONE.0211053

Simmons, B. I., Cirtwill, A. R., Baker, N. J., Wauchope, H. S., Dicks, L. V., Stouffer, D. B., & Sutherland, W. J. (2019). Motifs in bipartite ecological networks: uncovering indirect interactions. Oikos, 128(2), 154–170. https://doi.org/10.1111/OIK.05670

Siugzdaite, R., Bathelt, J., Holmes, J., & Astle, D. E. (2020). Transdiagnostic Brain Mapping in Developmental Disorders. Current Biology : CB, 30(7), 1245-1257.e4. https://doi.org/10.1016/J.CUB.2020.01.078

Sokolova, E., Groot, P., Claassen, T., Van Hulzen, K. J., Glennon, J. C., Franke, B., Heskes, T., & Buitelaar, J. (2016). Statistical Evidence Suggests that Inattention Drives Hyperactivity/Impulsivity in Attention Deficit-Hyperactivity Disorder. PLOS ONE, 11(10), e0165120. https://doi.org/10.1371/JOURNAL.PONE.0165120

Sporns, O., Tononi, G., & Kötter, R. (2005). The human connectome: A structural description of the human brain. PLoS Computational Biology, 1(4), 0245–0251. https://doi.org/10.1371/JOURNAL.PCBI.0010042

Stone, L. L., Janssens, J. M. A. M., Vermulst, A. A., Van Der Maten, M., Engels, R. C. M. E., & Otten, R. (2015). The Strengths and Difficulties Questionnaire: Psychometric properties of the parent and teacher version in children aged 4-7. BMC Psychology, 3(1), 1–12. https://doi.org/10.1186/S40359-015-0061-8

Suárez, L. E., Richards, B. A., Lajoie, G., & Misic, B. (2021). Learning function from structure in neuromorphic networks. Nature Machine Intelligence 2021 3:9, 3(9), 771–786. https://doi.org/10.1038/s42256-021-00376-1

Talkovsky, A. M., Green, K. L., Osegueda, A., & Norton, P. J. (2017). Secondary depression in transdiagnostic group cognitive behavioral therapy among individuals diagnosed with anxiety disorders. Journal of Anxiety Disorders, 46, 56–64. https://doi.org/10.1016/J.JANXDIS.2016.09.008

Theis, N., Rubin, J., Cape, J., Iyengar, S., Gur, R. E., Gur, R. C., Roalf, D. R., Pogue-Geile, M. F., Almasy, L., Nimgaonkar, V. L., & Prasad, K. M. (2021). Evaluating Network Threshold Selection for Structural and Functional Brain Connectomes. BioRxiv, 2021.10.09.463759. https://doi.org/10.1101/2021.10.09.463759

Tokuda, T., Yamashita, O., & Yoshimoto, J. (2021). Multiple clustering for identifying subject clusters and brain sub-networks using functional connectivity matrices without vectorization. Neural Networks, 142, 269–287. https://doi.org/10.1016/J.NEUNET.2021.05.016

Toplak, M. E., Bucciarelli, S. M., Jain, U., & Tannock, R. (2009). Executive functions: performance-based measures and the behavior rating inventory of executive function (BRIEF) in adolescents with attention deficit/hyperactivity disorder (ADHD). Child Neuropsychology, 15(1), 53–72. https://doi.org/10.1080/09297040802070929

Tournier, J. D., Smith, R., Raffelt, D., Tabbara, R., Dhollander, T., Pietsch, M., Christiaens, D., Jeurissen, B., Yeh, C. H., & Connelly, A. (2019). MRtrix3: A fast, flexible and open software framework for medical image processing and visualisation. NeuroImage, 202, 116137. https://doi.org/10.1016/J.NEUROIMAGE.2019.116137

Trudeau, R. J. (2013). Introduction to graph theory (2nd ed.). Courier Corporation.

Van Essen, D. C., & Barch, D. M. (2015). The human connectome in health and psychopathology. World Psychiatry : Official Journal of the World Psychiatric Association (WPA), 14(2), 154–157. https://doi.org/10.1002/WPS.20228

van Wijk, B. C. M., Stam, C. J., & Daffertshofer, A. (2010). Comparing Brain Networks of Different Size and Connectivity Density Using Graph Theory. PLOS ONE, 5(10), e13701. https://doi.org/10.1371/JOURNAL.PONE.0013701

Vanden Bussche, A. B., Haug, N. A., Ball, T. M., Padula, C. B., Goldstein-Pierarski, A. N., & Williams, L. M. (2017). Utilizing a transdiagnostic neuroscience-informed approach to differentiate the components of a complex clinical presentation: A case report. Personalized Medicine in Psychiatry, 3, 30–37. https://doi.org/10.1016/J.PMIP.2017.04.001

Viola, A., Balsamo, L., Neglia, J. P., Brouwers, P., Ma, X., & Kadan-Lottick, N. S. (2017). The Behavior Rating Inventory of Executive Function (BRIEF) to Identify Pediatric Acute Lymphoblastic Leukemia (ALL) Survivors At Risk for Neurocognitive Impairment. Journal of Pediatric Hematology/Oncology, 39(3), 174. https://doi.org/10.1097/MPH.0000000000000761

Wang, L., Zhu, C., He, Y., Zang, Y., Cao, Q., Zhang, H., Zhong, Q., & Wang, Y. (2009). Altered small-world brain functional networks in children with attention-deficit/hyperactivity disorder. Human Brain Mapping, 30(2), 638–649. https://doi.org/10.1002/HBM.20530

Wang, Y., Zuo, C., Xu, Q., Liao, S., Kanji, M., & Wang, D. (2020). Altered resting functional network topology assessed using graph theory in youth with attention-deficit/hyperactivity disorder. Progress in Neuro-Psychopharmacology and Biological Psychiatry, 98, 109796. https://doi.org/10.1016/J.PNPBP.2019.109796

Watts, D. J., & Strogatz, S. H. (1998). Collective dynamics of ‘small-world’ networks. Nature 1998 393:6684, 393(6684), 440–442. https://doi.org/10.1038/30918

Wechsler, D. (2003). Wechsler Intelligence Scale for Children–Fourth Edition (WISC-IV). San Antonio, TX: NCS Pearson.

Wilkinson, M., Wang, R., van der Kouwe, A., & Takahashi, E. (2016). White and gray matter fiber pathways in autism spectrum disorder revealed by ex vivo diffusion MR tractography. Brain and Behavior, 6(7), e00483. https://doi.org/10.1002/BRB3.483

Wold, H. (1975). Path Models with Latent Variables: The NIPALS Approach. Quantitative Sociology, 307–357. https://doi.org/10.1016/B978-0-12-103950-9.50017-4

Wold, H. (2004). Partial Least Squares. Encyclopedia of Statistical Sciences. https://doi.org/10.1002/0471667196.ESS1914

Xia M, Wang J, He Y (2013) BrainNet Viewer: A Network Visualization Tool for Human Brain Connectomics. PLoS ONE, 8, e68910.

Yang, Y. Y., Mahfouf, M., & Panoutsos, G. (2012). Probabilistic characterisation of model error using Gaussian mixture model—With application to Charpy impact energy prediction for alloy steel. Control Engineering Practice, 20(1), 82–92. https://doi.org/10.1016/J.CONENGPRAC.2011.10.001

Yendiki, A., Koldewyn, K., Kakunoori, S., Kanwisher, N., & Fischl, B. (2014). Spurious group differences due to head motion in a diffusion MRI study. NeuroImage, 88, 79–90. https://doi.org/10.1016/j.neuroimage.2013.11.027

Yufik, Y. M., Sengupta, B., & Friston, K. (2017). Editorial: Self-organization in the nervous system. Frontiers in Systems Neuroscience, 11, 69. https://doi.org/10.3389/FNSYS.2017.00069/BIBTEX

Zarrabi, M., Shahrivar, Z., Doost, M. T., Khademi, M., & Nejad, G. Z. (2015). Concurrent Validity of the Behavior Rating Inventory of Executive Function in Children With Attention Deficit Hyperactivity Disorder. Iranian Journal of Psychiatry and Behavioral Sciences, 9(1), 213. https://doi.org/10.17795/IJPBS213

Zhang-James, Y., Helminen, E. C., Liu, J., Busatto, G. F., Calvo, A., Cercignani, M., Chaim-Avancini, T. M., Gabel, M. C., Harrison, N. A., Lazaro, L., Lera-Miguel, S., Louza, M. R., Nicolau, R., Rosa, P. G. P., Schulte-Rutte, M., Zanetti, M. V., Ambrosino, S., Asherson, P., Banaschewski, T., … Faraone, S. V. (2021). Evidence for similar structural brain anomalies in youth and adult attention-deficit/hyperactivity disorder: a machine learning analysis. Translational Psychiatry 2021 11:1, 11(1), 1–9. https://doi.org/10.1038/s41398-021-01201-4

Ziegler, J. C., Perry, C., & Zorzi, M. (2020). Learning to Read and Dyslexia: From Theory to Intervention Through Personalized Computational Models. Current Directions in Psychological Science, 29(3), 293–300. https://doi.org/10.1177/0963721420915873

