## Supplement for "Exploring Neural Heterogeneity in Inattention and Hyperactivity"

***Supplementary Figure 1:*** *Scree plot displaying eigenvalues corresponding to n-factor solutions from our Exploratory Factor Analysis.*

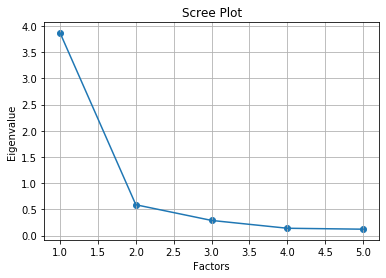

***Supplementary Figure 2:*** *Schematic representation of Partial Least Squares (PLS) Regression analysis used to compare inattention/hyperactivity factor scores and nodal degree, clustering coefficient, and communicability values across the entire sample (n = 383).*

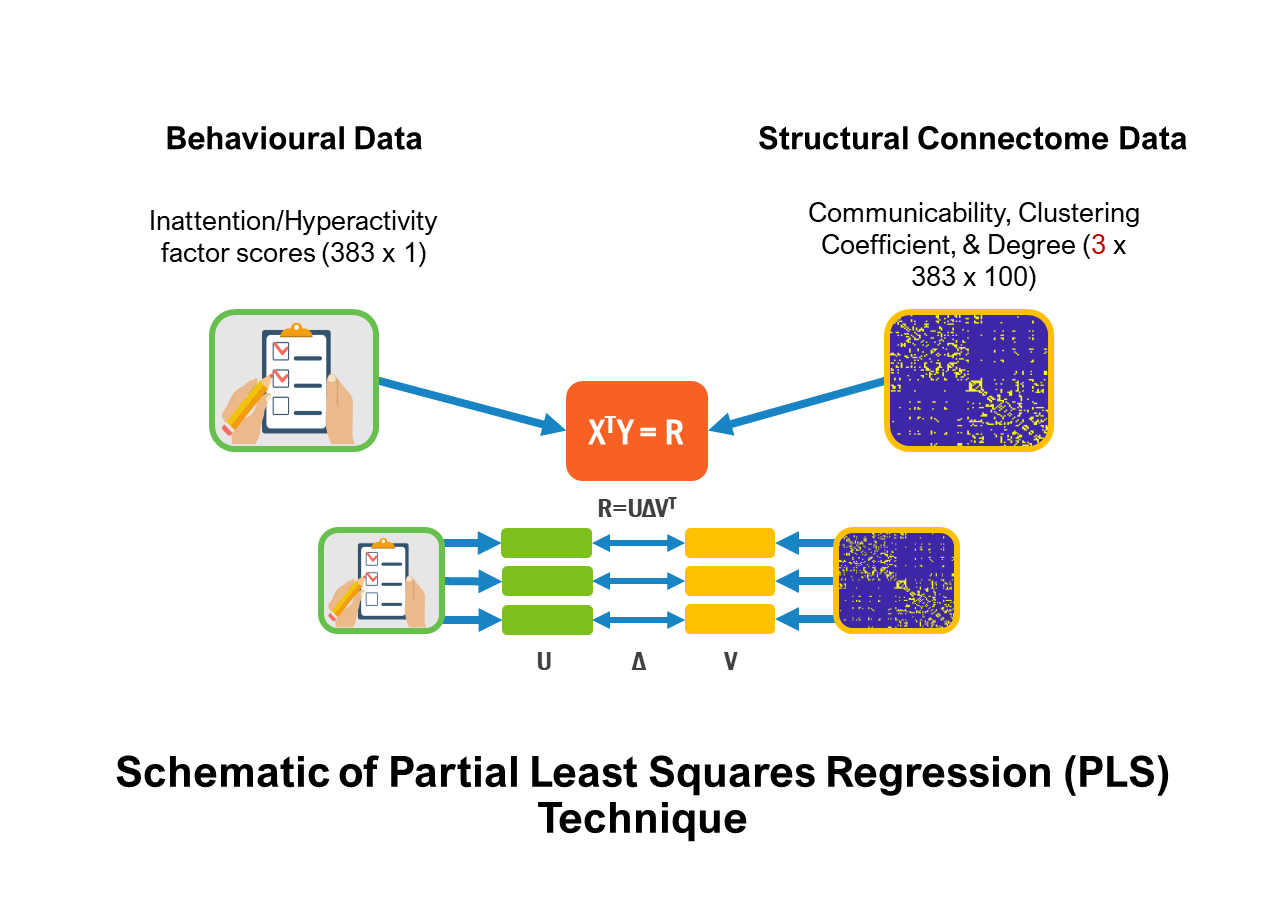

***Supplementary Table 1:*** *Silhouette values for n-cluster solutions from k-means clustering performed on the entire sample (n = 383). Multivariate outliers were identified using MATLAB’s robustcov function. After these outliers were excluded, nodewise graph measure data from n = 379 participants were included in the k-means clustering procedure.*

| ***Number of Clusters*** | ***Silhouette Coefficient*** | ***Cluster sizes*** |
| --- | --- | --- |
| **2** | 0.6028 | n = 216, 163 |
| **3** | 0.5673 | n = 141, 139, 99 |
| **4** | 0.5262 | n = 98, 70, 100, 111 |
| **5** | 0.4745 | n = 64, 70, 81, 93, 71 |
| **6** | 0.5031 | n = 70, 77, 63, 85, 25, 59 |
| **7** | 0.5032 | n = 65, 56, 24, 66, 59, 70, 39 |
| **8** | 0.4719 | n = 40, 64, 63, 75, 28, 56, 28, 25 |
| **9** | 0.5141 | n = 61, 26, 15, 47, 42, 42, 51, 53, 42 |

***Supplementary Table 2****: Silhouette values for alternative n-cluster solutions for inattentive/hyperactive sample (n = 232).*

| ***Number of Clusters*** | ***Silhouette Coefficient*** | ***Cluster sizes*** |
| --- | --- | --- |
| **2** | **0.6808** | **n = 109, 123** |
| **3** | 0.5860 | n = 89, 84, 59 |
| **4** | 0.5175 | n = 67, 90, 45, 30 |
| **5** | 0.4716 | n = 44, 63, 45, 37, 43 |
| **6** | 0.4699 | n = 42, 38, 39, 40, 39, 34 |
| **7** | 0.5167 | n = 35, 15, 38, 27, 41, 50, 26 |
| **8** | 0.4594 | n = 19, 32, 42, 30, 30, 32, 17, 30 |
| **9** | 0.4997 | n = 18, 29, 11, 48, 33, 25, 25, 35, 8 |

***Supplementary Table 3:*** *Behavioural (in blue) and cognitive (in red) characteristics of inattentive/hyperactive children (Conners Questionnaire scores >60; n = 232) compared to children without elevated inattention and hyperactivity (Conners Questionnaire scores <60; n = 151), in addition to the full sample (n = 383). Values are displayed in ‘Mean (SD)’ format. ANOVAs were performed to compare the three groups on 19 measures of behaviour and cognition.*

| ***Measure*** | ***Total sample***  ***(n = 383)*** | ***I/H subsample***  ***(n = 232)*** | ***Discarded sample***  ***(n = 151)*** | ***F*** | ***p*** |
| --- | --- | --- | --- | --- | --- |
| **SDQ (Total)** | 15.87 *(8.66)* | 20.30 *(6.92)* | 6.14 *(5.21)* | 95.432 | **1.148 x 10^-37^** |
| **SDQ (Emotion Regulation)** | 3.71 *(2.84)* | 4.48 *(2.80)* | 1.87 *(2.15)* | 22.863 | **2.294 x 10^-10^** |
| **SDQ (Conduct)** | 2.80 *(2.38)* | 3.73 *(2.38)* | 1.14 *(1.35)* | 50.060 | **3.871 x 10^-21^** |
| **SDQ (Hyperactivity)** | 6.49 *(3.19)* | 8.49 *(1.72)* | 1.96 *(1.93)* | 163.207 | **1.313 x 10^-59^** |
| **SDQ (Peer Problems)** | 2.87 *(2.67)* | 3.61 (2.70) | 1.18 *(1.58)* | 23.443 | **1.329 x 10^-10^** |
| **SDQ (Prosocial)** | 7.31 *(2.22)* | 6.80 *(2.22)* | 8.61 *(1.71)* | 16.753 | **7.609 x 10^-08^** |
| **Conners (Inattention)** | 75.00 *(16.30)* | 83.82 *(7.46)* | 47.73 *(6.58)* | 114.137 | **5.401 x 10^-44^** |
| **Conners (Hyperactivity/Impulsivity)** | 69.71 *(17.23)* | 81.38 *(9.64)* | 48.21 *(5.85)* | 212.225 | **8.005 x 10^-74^** |
| **Conners (Learning Problems)** | 70.31 *(15.91)* | 75.96 *(12.21)* | 52.79 *(13.11)* | 41.0102 | **1.243 x 10^-17^** |
| **Conners (Executive Function)** | 70.27 *(15.70)* | 77.58 *(10.45)* | 48.57 *(9.04)* | 77.094 | **3.587 x 10^-31^** |
| **Conners (Aggression)** | 59.85 *(16.48)* | 65.61 *(17.21)* | 49.25 *(7.38)* | 40.432 | **2.104 x 10^-17^** |
| **Conners (Peer Relations)** | 68.91 *(18.67)* | 74.96 *(17.36)* | 53.72 *(13.22)* | 32.527 | **2.863 x 10^-14^** |
| **WASI-II** | 46.01 *(10.14)* | 44.23 *(9.46)* | 51.42 *(10.52)* | 10.249 | **4.050 x 10^-5^** |
| **Peabody Picture Vocabulary Test** | 93.65 *(16.67)* | 92.68 *(9.37)* | 109.81 *(16.65)* | 0.724 | 0.485 |
| **PhAB (Alliteration)** | 103.59 *(9.50)* | 101.64 *(16.21)* | 97.26 *(8.22)* | 10.842 | **2.278 x 10^-5^** |
| **AWMA (Digit Recall)** | 96.58 *(17.28)* | 88.75 *(14.00)* | 104.49 *(17.02)* | 8.921 | **0.0001** |
| **AWMA (Dot Matrix)** | 94.60 *(16.23)* | 93.18 *(15.98)* | 102.75 *(15.22)* | 4.163 | **0.0159** |
| **AWMA (Digit Back)** | 95.77 *(14.30)* | 91.63 *(15.88)* | 104.68 *(14.61)* | 18.174 | **1.963 x 10^-8^** |
| **AWMA (Mr X)** | 99.85 *(16.06)* | 92.75 *(12.86)* | 105.71 *(16.67)* | 13.049 | **2.685 x 10^-6^** |

***Supplementary Figure 3****: Schematic representation of General Linear Model (GLM) analyses comparing behavioural questionnaire scores between the inattentive/hyperactive clusters.*

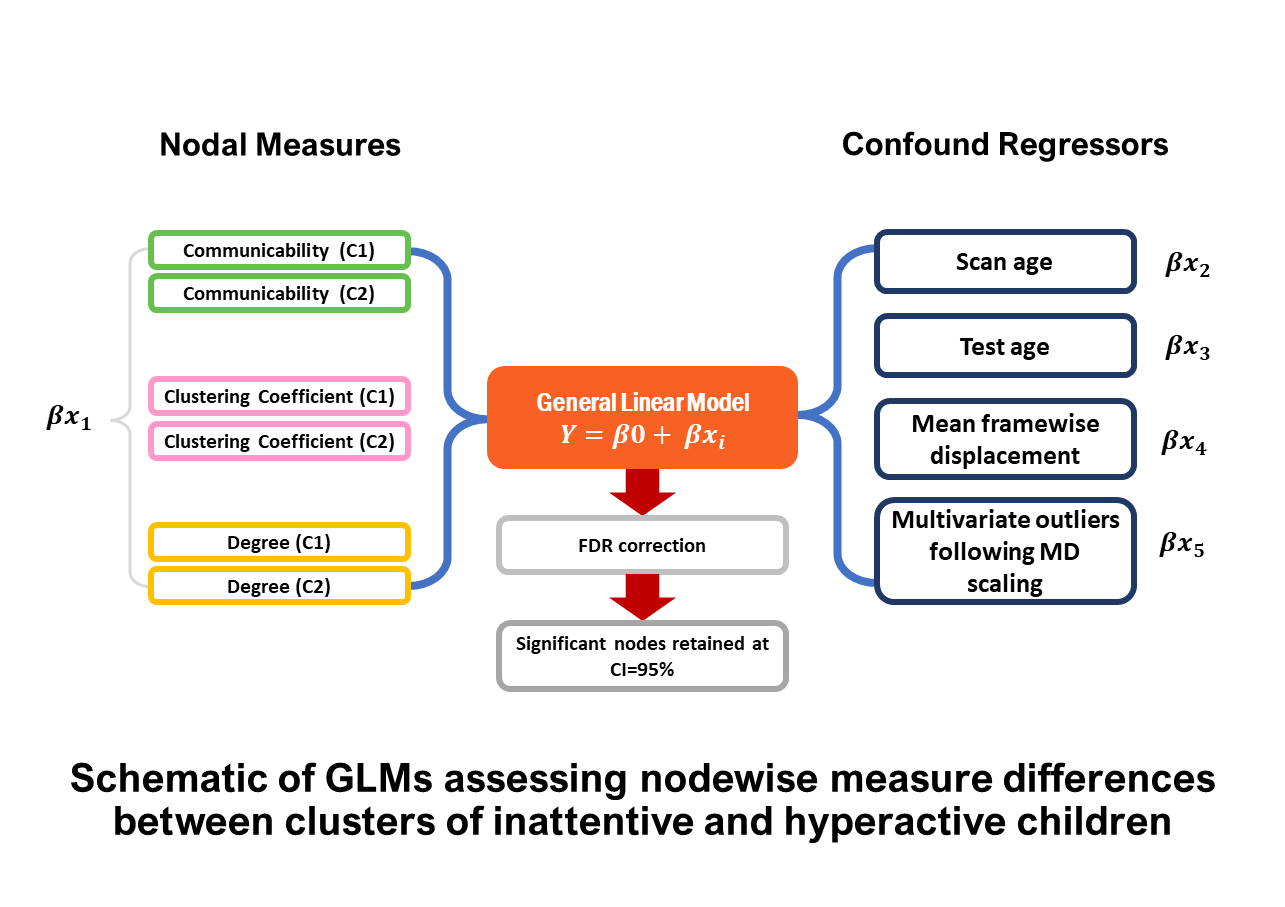

***Supplementary Figure 4****: Schematic representation of General Linear Model (GLM) analyses comparing behavioural questionnaire scores between the inattentive/hyperactive clusters.*

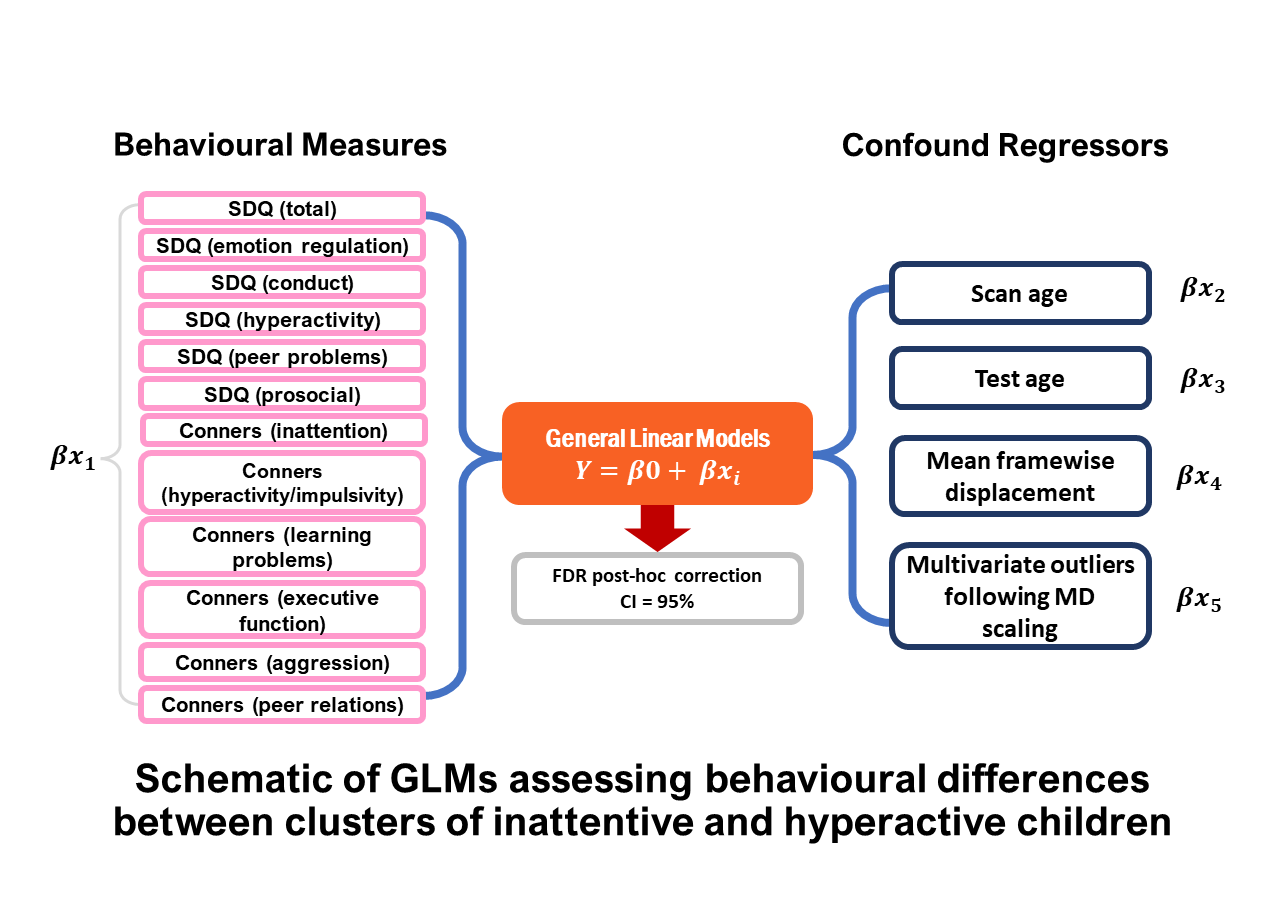

***Supplementary Figure*** ***5****: Schematic representation of General Linear Model (GLM) analyses comparing cognitive test scores between the inattentive/hyperactive clusters.*

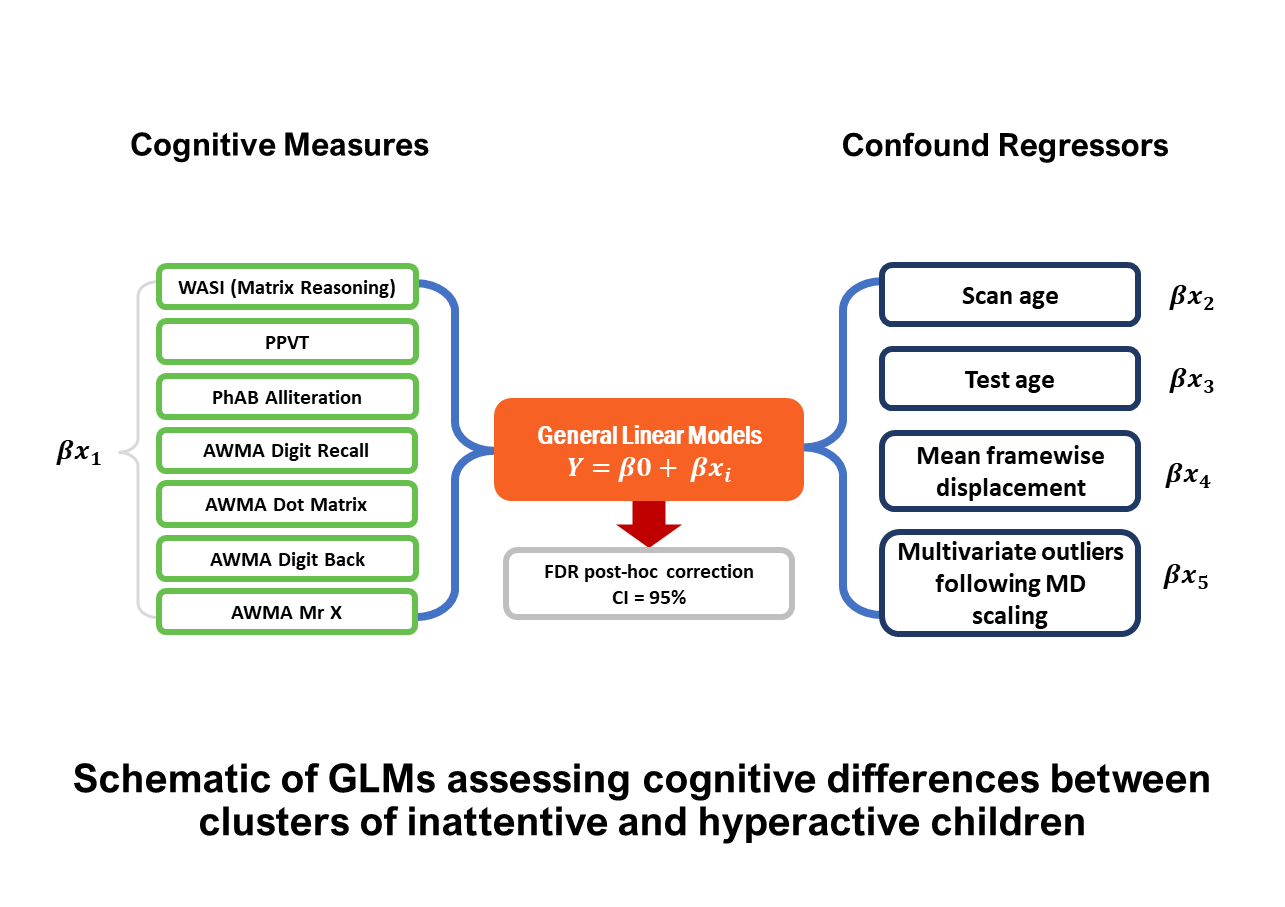

***Supplementary Figure 6:*** *Histograms displaying distributions of confound regressor values (mean framewise displacement and age) across the inattentive/hyperactive sample (n = 232).*

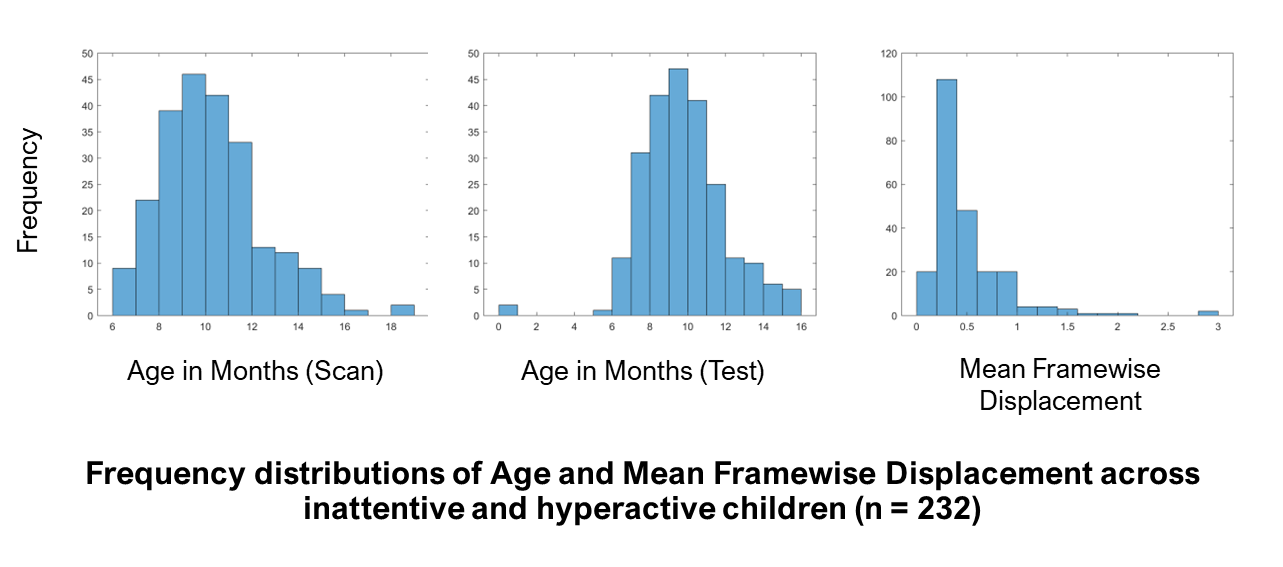
